# Simulating BRAFV600E-MEK-ERK signalling dynamics in response to vertical inhibition treatment strategies

**DOI:** 10.1101/2023.12.12.571169

**Authors:** Alice De Carli, Yury Kapelyukh, Jochen Kursawe, Mark A.J. Chaplain, C. Roland Wolf, Sara Hamis

## Abstract

In vertical inhibition treatment strategies, multiple components of an intracellular pathway are simultaneously inhibited. Vertical inhibition of the BRAFV600E-MEK-ERK signalling pathway is a standard of care for treating BRAFV600E-mutated melanoma where two targeted cancer drugs, a BRAFV600E-inhibitor, and a MEK-inhibitor, are administered in combination. Targeted therapies have been linked to early onsets of drug resistance, and thus treatment strategies of higher complexities and lower doses have been proposed as alternatives to current clinical strategies. However, finding optimal complex, low-dose treatment strategies is a challenge, as it is possible to design more treatment strategies than are feasibly testable in experimental settings.

To quantitatively address this challenge, we develop a mathematical model of BRAFV600E-MEK-ERK signalling dynamics in response to combinations of the BRAFV600E-inhibitor dabrafenib (DBF), the MEK-inhibitor trametinib (TMT), and the ERK-inhibitor SCH772984 (SCH). From a model of the BRAFV600E-MEK-ERK pathway, and a set of molecular-level drug-protein interactions, we extract a system of chemical reactions that is parameterized by in vitro data and converted to a system of ordinary differential equations (ODEs) using the law of mass action. The ODEs are solved numerically to produce simulations of how pathway-component concentrations change over time in response to different treatment strategies, *i*.*e*., inhibitor combinations and doses. The model can thus be used to limit the search space for effective treatment strategies that target the BRAFV600E-MEK-ERK pathway and warrant further experimental investigation. The results demonstrate that DBF and DBF-TMT-SCH therapies show marked sensitivity to BRAFV600E concentrations in silico, whilst TMT and SCH monotherapies do not.

## Introduction

Cancer cells have properties that differentiate them from normal cells of the human body. Targeted cancer drugs work by specifically targeting those properties.^1^ The clinical use of small-molecule targeted cancer drugs, which began in the early 2000s, has improved survival rates for multiple cancers.^2^ However, targeted cancer drugs have been linked to early onsets of drug resistance in patients,^3,4,5^ which poses a major clinical question: How do we avoid or delay resistance to targeted cancer drugs? Proposed strategies to approach this question include altering (i) drug combinations, (ii) drug doses, and (iii) drug schedules. By altering (i-iii), it is possible to design more treatment strategies than can be feasibly tested. Therefore, optimal strategies cannot be found by pre-clinical experiments and clinical trials alone. To limit the search space for effective treatment strategies that warrant pre-clinical and clinical investigation, rational and quantitative approaches can be used. Rational treatment strategies refer to strategies informed by mechanistic understandings of how treatments target cancer cells’ abilities to survive and proliferate.^4,6^ One such rational treatment strategy is vertical inhibition, where multiple components of an intracellular signalling pathway are simultaneously inhibited.^5^

For treating BRAFV600E-mutated melanoma, for instance, vertical inhibition is now a standard of care.^7^ The BRAFV600E mutation causes hyper-activated signalling in the intracellular BRAFV600E-MEK-ERK pathway, in which BRAFV600E activates MEK via phosphorylation events, which in turn activates ERK via phosphorylation events.^8^ Activated ERK promotes cell survival and cell proliferation by activating numerous substrates in the cell nucleus and cytoplasm. Thus, hyper-activated BRAFV600E-MEK-ERK signalling may result in high ERK activity and, by extension, uncontrolled cell proliferation.^9,10^ ERK-activated substrates include structural proteins, nuclear envelope proteins and transcription factors such as the cancer-associated substrate c-Myc.^11^ c-Myc is involved in regulating multiple cell functions, including cell cycle progression, cell differentiation and cell death.^12^ In normal cells, c-Myc expression is tightly controlled. In cancer cells, however, c-Myc expression has been observed to be dysregulated, entailing increased c-Myc concentration and uncontrolled cell proliferation.^13,14^ This overexpression is promoted by upstream signalling of *e*.*g*. the BRAFV600E-MEK-ERK pathway.^12^

Vertical inhibition of the BRAFV600E-MEK-ERK pathway ultimately acts to suppress ERK’s ability to activate substrates. The rationale is that by inhibiting not only the BRAFV600E oncogene but also downstream pathway components such as MEK and ERK, treatments should be more effective and drug resistance should be impeded. In support of this rationale, previous experimental results indicate that when multiple components in a signalling pathway are inhibited, the risk for cancer cells evading drug effects via adaptive feedback reactivation of pathway signalling is reduced.^15,16,17,18^ Moreover, experimental results show that high-dose monotherapies of inhibitors that target the BRAFV600E-MEK-ERK pathway exacerbate the onset of drug resistance in vivo, by creating evolutionary pressures that select for drug-resistant cell clones.^19^ These evolutionary pressures decrease with increased treatment complexities, enabled via vertical inhibition.^19^ The risks of high-dose drug treatments are important to note, as cancer drugs are commonly clinically administered at doses that are determined in the early, dose-escalation stages of clinical trials. This is the case for the BRAFV600E-specific inhibitor dabrafenib (DBF), and the MEK-inhibitor trametinib (TMT) which are administered together for treating BRAFV600E-mutated melanoma.^20,21,22^ Low-dose vertical inhibition strategies have been shown to provide promising alternatives to high-dose strategies in vitro and in vivo.^23^

As a complement to experimental and clinical studies, mathematical models can be used to simulate and analyze pathway signalling dynamics^24,25,26,27,28,29^ and quantitatively identify vertical inhibition, low-dose drug combinations that yield synergistic treatment responses in silico.^30^ Notably, in 1996, Huang and Ferrell developed a reaction network model that describes signalling dynamics in general MAPKKK-MAPKK-MAPK cascades (MAPK cascades),^25^ of which the BRAFV600E-MEK-ERK pathway is one. Whilst Huang and Ferrell predicted ultrasensitive dynamics in MAPK cascades, such that ERK goes from being non-activated to activated in a switch-like manner, later work by Aoki et al. showed that ERK phosphorylation in HeLa cells is graded or, in other words, processive.^31^ The authors also suggested that, under physiological conditions where molecular crowding occurs, ERK phosphorylation is quasi-processive. Building on observations from yeast experiments, Takahashi and Pryciak concluded that the MAPK cascade is inherently ultrasensitive, but that the pathway can be converted to a processive form via interactions with scaffold proteins.^32^ Scaffold proteins are, indeed, crucial for signal regulation and activation in MAPK cascades. This was emphasised by Levchenko et al. who developed a computational model in which MAPK cascade components interact with a generic scaffold protein. Model analysis revealed that scaffold protein concentrations significantly alter cascade signalling dynamics.^33^ The importance of cascade-scaffold interactions was also marked by Tian et al. who studied Ras signalling nanoclusters, *i*.*e*., discrete plasma membrane nanodomains. The authors showed that spatial Ras organisation crucially impacts pathway signalling.^34^ Furthermore, to investigate downstream effects of ERK signalling, Lee et al. developed a mathematical model that integrates signalling dynamics in the ERK and the phosphatidylinositol 3-kinase (PI3K) pathways, the two effector pathways that regulate Myc.^35^ Their model suggested that Myc stability is sensitive to temporal aspects of ERK and PI3K signalling. Building on Huang and Ferrell’s modelling work,^25^ we (some of the authors) recently developed a mathematical model and a computational framework for simulating signalling dynamics in the BRAFV600E-MEK-ERK pathway in response to vertical inhibition with two drugs: DBF and TMT.^30^ Our study focused on signalling dynamics in response to pathway-drug interactions and therefore omitted details pertaining to *e*.*g*., scaffold proteins, spatial molecular structures, and other intracellular pathways. In the previous study, we calculated ERK activities in response to varying concentrations of drugs, adenosine triphosphate (ATP), and pathway components. The model provided a mechanistic understanding on a molecular level of when DBF-TMT synergy occurs and identified intracellular BRAFV600E and ATP levels as factors in drug resistance.

Beyond two-component vertical inhibition strategies, combining BRAFV600E, MEK, and ERK inhibitors has shown promising results in pre-clinical experiments.^19^ Therefore, ERK-inhibitors have been proposed to be used in combination with DBF and TMT, as part of a three-drug vertical inhibition strategy.^19^ To investigate such strategies in silico we here extend our previous mathematical model^30^ to also include an ATP-competitive ERK-inhibitor. The model is parameterized by data pertaining specifically to the ERK-inhibitor SCH772984,^36^ but the parameter values can be adjusted to study other ATP-competitive ERK-inhibitors. We use the model to simulate BRAFV600E-MEK-ERK signalling dynamics in response to various combinations and doses of BRAFV600E, MEK, and ERK inhibitors. Our results also identify two and three-component vertical inhibition dose combinations that yield synergistic treatment responses in silico, in terms of inhibiting the activation of an ERK-activated substrate.

## Results

We simulate signalling dynamics in the BRAFV600E-MEK-ERK pathway using a mathematical model and a computational framework.^30^ The mathematical model comprises an abstraction of the intracellular BRAFV600E-MEK-ERK pathway, and a set of possible interactions that can occur between pathway components and inhibitors. In order to evaluate the effect of treatment strategies that involve ERK-inhibitors, we extend the pathway model to include the ERK-activated substrate c-Myc, which we refer to as S, downstream of ERK. From the c-Myc extended pathway model and the set of possible interactions, we extract a system of chemical reactions that describe pathway signalling in response to vertical inhibition. Using our previously developed computational framework^30^ that builds on the law of mass action, we convert the system of chemical reactions into a system of ordinary differential equations (ODEs) that are solved numerically to produce simulations of pathway signalling dynamics. These simulations show how concentrations of the pathway components change over time, in response to different inhibitor combinations and concentrations. We choose phosphorylated c-Myc (pS) to be our main model output of interest, and the simulated treatments ultimately act to suppress the amount of pS in the modelled system. By altering the initial inhibitor concentrations, we simulate treatment responses to vertical inhibition strategies that target one, two, or three components of the pathway. To simulate pathway dynamics in the absence of a specific inhibitor, we simply set the initial concentration of that inhibitor to zero. Throughout this study, the total amount of inhibitor concentrations is constant over the time courses of the simulations.

### Simulating treatment responses to monotherapies that inhibit the BRAFV600E-MEK-ERK pathway

We first simulate pathway dynamics in response to monotherapies that inhibit BRAFV600E, MEK, or ERK activity. Our main simulation output of interest is *S*_*act*_(*t*), the fraction of simultaneously free and activated substrate S in the system (Eq. 1, Methods). We are interested in investigating how this model output changes in time, in response to different doses of the BRAFV600E-inhibitor DBF, the MEK-inhibitor TMT, or the ERK-inhibitor SCH. We are also interested in understanding how intracellular concentrations of BRAFV600E and ATP impact treatment responses in silico. We choose an initial condition in which only free and non-phosphorylated MEK, ERK, and S are present in the system, in line with previous studies^25,30^ (Supplementary Material, Supplementary Note 3.2). Hence the substrate S is not immediately activated when the simulations start. Instead, S activation occurs after MEK and ERK activation, as regulated by the pathway dynamics. This transient behaviour of S activation is illustrated in Fig. 1, where monotherapy responses to DBF, TMT, and SCH are plotted over increasing inhibitor concentrations for six different simulation time points.

**Fig. 1:**
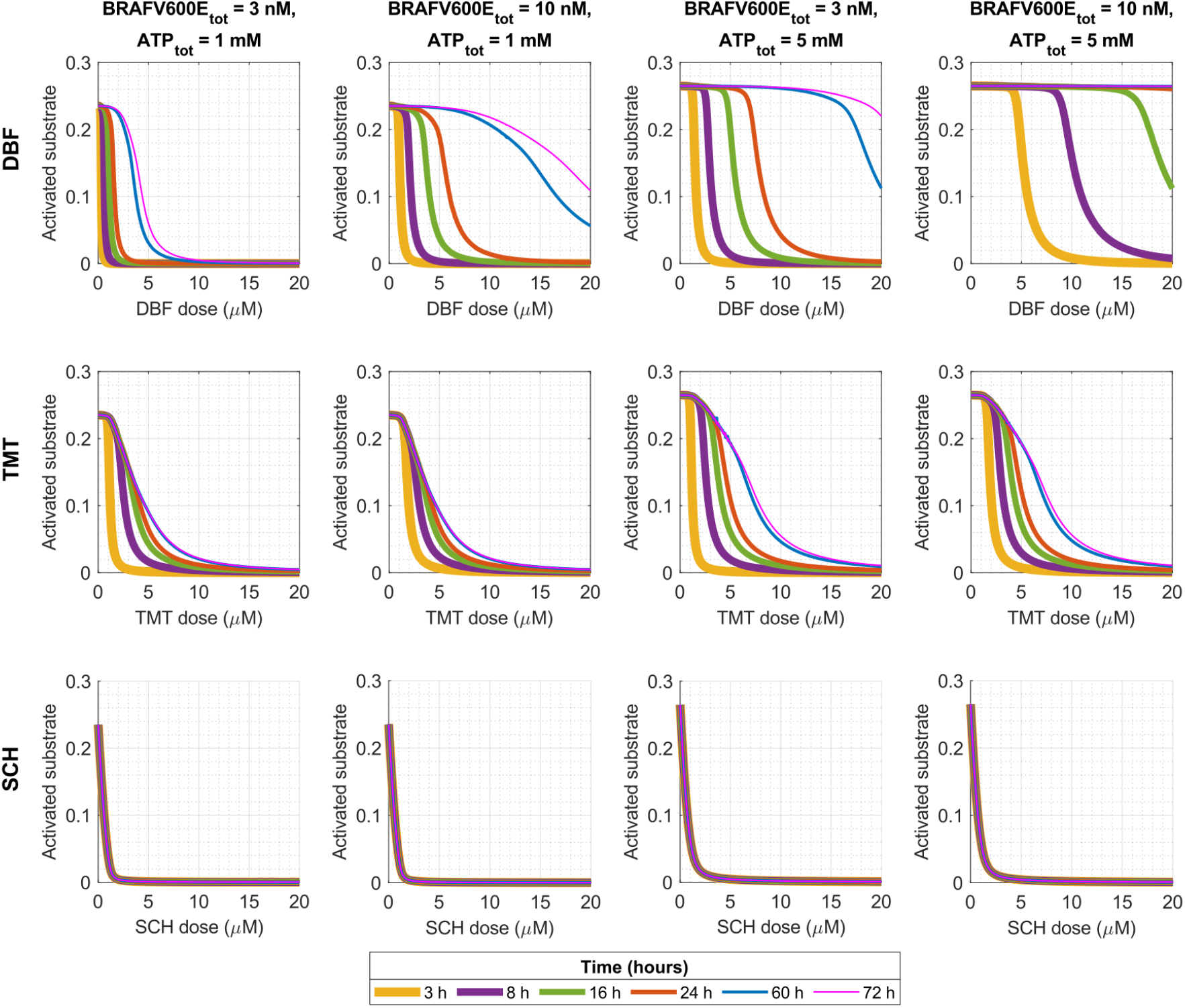
Mathematical modelling quantifies in silico treatment responses to monotherapies that target components of the BRAFV600E-MEK-ERK pathway. The plots show activated substrate levels (*S*_*act*_(*t*), Eq. 1, Methods) over time in response to different doses of the BRAFV600E-inhibitor DBF (top row), the MEK-inhibitor TMT (middle row), and the ERK-inhibitor SCH (bottom row). The plots show activated substrate levels at six simulation time points (3, 8, 16, 24, 60 and 72 hours). The total BRAFV600E and ATP concentrations vary between the columns, where baseline values (BRAFV600E_tot_=3 nM, ATP_tot_=1 mM) are used in the left-most column

With the baseline values of total BRAFV600E and ATP concentrations, which are estimated to correspond to physiological values,^30^ TMT and SCH monotherapies yield steady-state activated substrate levels within 72 hours for all doses (Fig. 1, left column). These steady-state levels are reached within 60 hours for TMT treatments, and within three hours for SCH treatments. In contrast, no steady-states are reached in response to DBF monotherapies. By comparing the two left-most columns in Fig. 1, we can study what happens to substrate activation dynamics when the total BRAFV600E concentration is increased from the 3 nM baseline value to 10 nM. This comparison shows that the increased BRAFV600E concentration raises the dose-response curves and thus desensitizes the pathway to DBF monotherapies. Furthermore, the simulation results show that the steady-state treatment responses to TMT and SCH monotherapies are not affected by the change in BRAFV600E concentrations from 3 nM to 10 nM, although the time durations to reach the steady-states do decrease with increasing BRAFV600E concentrations. This latter result is visible in Fig. 1 for TMT monotherapies at 3 hours. We note that the transient system behaviour of responses to SCH monotherapies is not visible from Fig. 1, as steady-state levels are reached within 3 hours. However, this transient system behaviour is observable from supplementary simulation results where we plot activated substrate levels over time (Supplementary Material, Supplementary Note 4).

Next, to investigate the role of ATP in treatment responses, we increase the total ATP concentration from the baseline value of 1 mM to 5 mM in the third column. We then compare the first and third columns in Fig. 1. Doing this, we see that the increase in ATP levels renders the pathway less sensitive to DBF, TMT, and SCH monotherapies at all regarded time points. These results follow from the fact that ATP is needed for the phosphorylation of MEK, ERK, and S. Hence, when intracellular ATP concentrations increase, so does the probability that ATP-binding sites on the catalyzing enzymes will be occupied by ATP molecules.

In summary, from the simulation results, we find that both BRAFV600E and ATP levels impact transient and absolute values of *S*_*act*_(*t*) in response to DBF monotherapies. Conversely, in response to TMT and SCH doses, ATP concentrations predominately impact the steady-state levels of *S*_*act*_(*t*), whilst BRAFV600E concentrations impact the time until steady-state levels are reached. This finding is substantiated by Fig. 1 (right), in which both total BRAFV600E and ATP concentrations have been elevated from the baseline values. Notably, the model predicts that DBF monotherapies are initially effective in suppressing substrate activation, but that this efficacy decreases over time. For instance, at 3 hours and baseline BRAFV600E and ATP concentrations, 0.45 *µ*M DBF suffices to achieve 90% inhibition of activated substrate 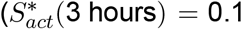, Eq. 3, Methods), whilst the same dose inhibits less than 1% substrate activation at 24 hours. On the other hand, a 1.15 *µ*M SCH dose achieves 90% inhibition of activated substrate for all time points at 3 hours and over. Even more strikingly, when the BRAFV600E concentration is increased to 10 nM, 90% inhibition of activated substrate is achieved by *e*.*g*., 1.57 *µ*M DBF at 3 hours, 8.71 *µ*M DBF at 24 hours, and 1.15 *µ*M SCH for time points at 3 hours and over. These results suggest that, due to (i) DBF-leakage, and (ii) DBF sensitivity to BRAFV600E concentrations, SCH monotherapies may be effective in complementing DBF therapies in inhibiting substrate activity.

### Simulating treatment responses to two-component vertical inhibition of the BRAFV600E-MEK-ERK pathway

We next simulate pathway dynamics in response to vertical inhibition treatment strategies that target two components of the BRAFV600E-MEK-ERK pathway. Activated substrate levels (*S*_*act*_, Eq. 1, Methods) in response to BRAFV600E and MEK inhibition (DBF-TMT), BRAFV600E and ERK inhibition (DBF-SCH), and MEK and ERK inhibition (TMT-SCH) are shown in the heatmaps for three selected time points (8, 16 and 24 hours) in Fig. 2. The time points are chosen to exemplify time windows of BRAFV600E-MEK-ERK signalling dynamics that are of interest from a treatment perspective, and are motivated by treatment schedules from the first-in-human study of DBF, where DBF was administered once, twice, or three times daily.^37^ The heatmaps demonstrate the dynamic behaviour of the system through the time-varying *S*_*act*_ levels. We use these heatmaps to visually identify dose combinations of two inhibitors that yield synergistic treatment responses in silico.

**Fig. 2:**
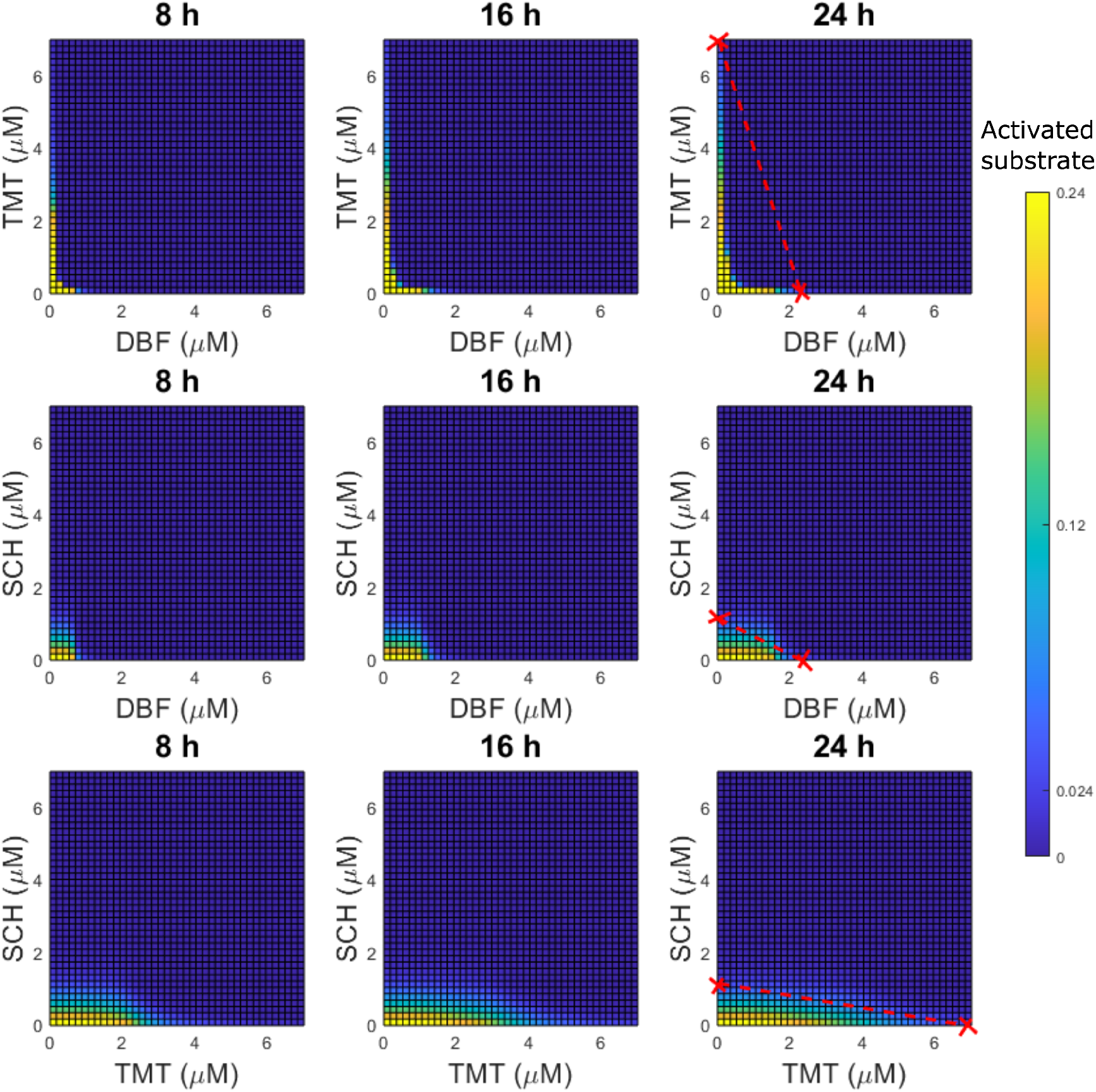
Mathematical modelling quantifies in silico treatment responses to vertical inhibition treatment strategies that target up to two components of the BRAFV600E-MEK-ERK pathway. The heatmaps show levels of substrate activity (*S*_*act*_(*t*), Eq. 1, Methods) in response to combination treatments of BRAFV600E and MEK inhibitors (DBF-TMT, top row), BRAFV600E and ERK inhibitors (DBF-SCH, middle row), and MEK and ERK inhibitors (TMT-SCH, bottom row) at three different simulation time points (8, 16 and 24 hours). Baseline values of total BRAFV600E (3 nM) and ATP (1 mM) concentrations are used in the simulations. The right-most plots include isoboles (lines) that connect the relevant monotherapy doses that yield 90% inhibition of substrate activity at 24 hours such that 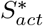 (24 hours) = 0.1, which corresponds to *S*_*act*_(24 hours) ≈ 0.024. Two-drug combination doses under the isoboles that achieve more than 90% activated substrate inhibition, such that 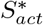 (24 hours) < 0.1, are here categorised as synergistic

In this study, we categorize a drug combination as being synergistic, or not, using the method of isoboles.^38^ To understand this method, consider an N-dimensional space where N denotes the number of drugs that we want to include in a drug combination. A drug dose combination can be describes as a point 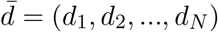 where *d*_*i*_ denotes the dose of drug *i*. In this space, we can create a synergy-isobole that connects all monotherapy doses that achieve some desired drug effect *E*^*∗*^. If a drug dose combination 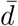 yields a drug effect that is higher than *E*^*∗*^, and 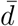 lies under the synergy-isobole, the drug combination is categorized as synergistic. Otherwise, the drug combination is categorized as non-synergistic. Here, we quantify the inhibitory effects of drugs in terms of 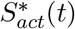 levels (Eq. 3, Methods), and in Fig. 2 we draw out isoboles between monotherapy doses that yield 90% inhibition at 24 hours, such that 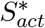 (24 hours) = 0.1. Thus, we categorize two-drug combination drug doses that achieve 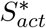 (24 hours) *<* 0.1 and lie under the isobole lines as synergistic. The 0.1 threshold value is chosen to reflect an inhibition level that we assume to be effective from a treatment perspective. This assumption is extrapolated from previous studies, where 80%,^39^ and 90%^40^ inhibition of ERK phosphorylation has been observed in patients with tumour regression. Importantly, the model can identify drug combinations that achieve any inhibition level of interest.

With the chosen synergy categorization, our results identify a broad range of synergistic DBF-TMT combinations (Fig. 2). These results are in line with our previous mathematical study,^30^ in which we identified DBF-TMT combinations that synergistically inhibit activated ERK levels (as opposed to activated substrate S levels which are the focal simulation output in the current study). However, our results do not identify synergistic DBF-SCH or TMT-SCH combinations (Fig. 2). This result is clearly visualised in the Supplementary Material (Supplementary Fig. 2) where we have re-plotted Fig. 2 with binary heatmaps that indicate if a treatment combination yields an inhibition of at least 90% or not. Note that these simulation results do not mean that DBF-SCH and TMT-SCH combinations are generally non-synergistic in practice, but rather that our pathway model alone cannot explain DBF-SCH or TMT-SCH synergies, according to our chosen synergy-categorization. We hypothesize that the inclusions of alternative pathway components or treatment-induced feedback loops in the pathway model might capture *eg*., potential DBF-SCH synergies. We also note that, from a treatment perspective, drug combinations need not be synergistic in order to be beneficial. Moreover, potential synergies might result from mechanisms at the cell population or tissue scale, rather than from pathway-intrinsic mechanisms at the intracellular scale. To capture such synergies mathematically, our intracellular model could, for example, be migrated to a cell population or tissue model via a multi-scale modelling approach.^41,42^

### Simulating treatment responses to three-component vertical inhibition of the BRAFV600E-MEK-ERK pathway

We, lastly, simulate BRAFV600E-MEK-ERK pathway dynamics in response to simultaneous BRAFV600E, MEK, and ERK inhibition. We are especially interested in identifying synergistic drug dose combinations of the BRAFV600E-inhibitor DBF, the MEK-inhibitor TMT, and the ERK-inhibitor SCH. Using the same isobole-based synergy categorization as in the previous subsection, a three-drug combination strategy is here classified as synergistic if the strategy achieves more than 90% inhibition of activated substrate at some time *t* (Eq. 3, Methods) and lies under a plane that connects the three monotherapy doses that yield 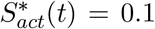. In Fig. 3, three-drug combination therapies that yield 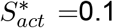 at 8, 16, and 24 hours are plotted on a surface. By picking any point on this surface, and using the axes to read out where in DBF-TMT-SCH space the regarded surface point lies, drug-dose combinations that yield 90% inhibition can be identified. To aid readability, we have included a numerical matrix of surface point drug-dose combinations in the Supplementary Material (Supplementary Fig. 3). Also included in Fig. 3 are synergy-isoboles for 90% inhibition. Thus, we can use the plots to graphically identify synergistic combinations of three drugs as any part of the 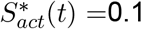 surface that lies under the synergy-isobole. Using this graphical method, we can observe a range of drug combinations that yield synergistic treatment responses. These follow from the synergistic in silico relationships that were observed for two-component vertical inhibition strategies in Fig. 2, and in our previous mathematical study.^30^ Note that the choice of using 90% inhibition in the drug-dose surfaces and synergy planes in Fig. 3 is guided by our assumption that 90% inhibition corresponds to an effective treatment objective, following our discussion in the previous subsection. In the Supplementary Material (Supplementary Note 6), we have repeated this graphical method to find drug combinations that achieve 25%, 50%, and 75% inhibition of substrate activity. More generally, the methodology used in this study can be used to find a range of one-, two-, and three-component vertical inhibition strategies that target the BRAFV600E-MEK-ERK pathway and achieve selected activity levels of any chosen pathway components.

**Fig. 3:**
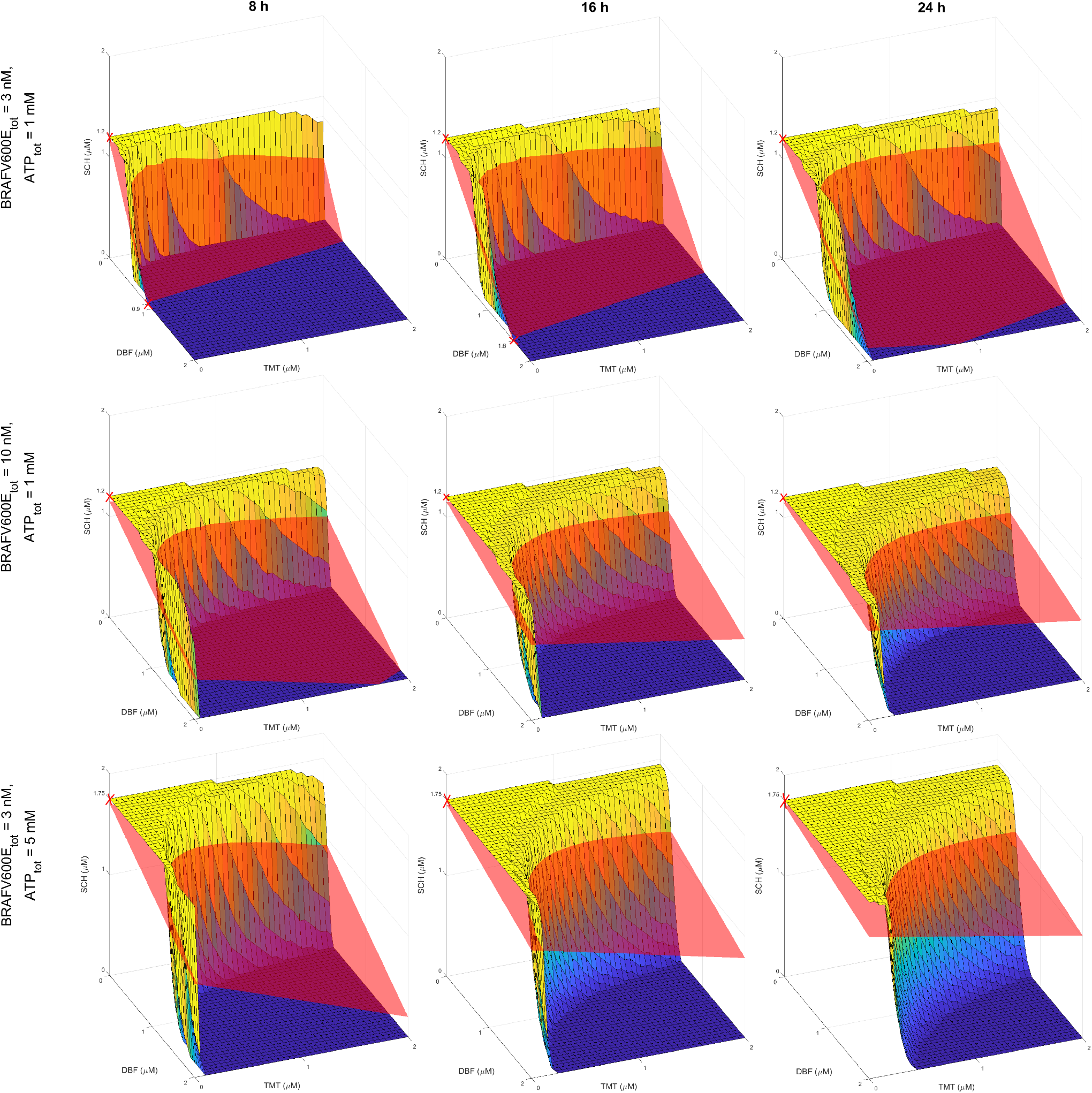
Mathematical modelling quantifies in silico treatment responses to vertical inhibition treatment strategies that target up to three components of the BRAFV600E-MEK-ERK pathway. The yellow-to-blue surface plots show DBF-TMT-SCH combination therapies that yield 90% substrate activity 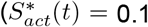, Eq. 3, Methods). The red synergy-isoboles (planes) connect the monotherapy doses that yield a 90% inhibition of substrate activity. Monotherapy doses that yield 90% substrate activity are marked with crosses. Treatment combinations, *i*.*e*., parts of the 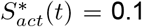 surfaces, that lie under the synergy-isoboles are here categorised as synergistic. Results are plotted at 8 (left column), 16 (middle column) and 24 (right column) hours for baseline, total ATP, and BRAFV600E concentrations (top row), and for elevated BRAFV600E concentrations (middle row) and ATP concentrations (bottom row)

The simulation results in Fig. 3 show that both 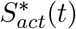 levels and the synergy-isoboles change over time and are sensitive to the total amount of BRAFV600E and ATP concentrations in the system.

Increasing the total BRAFV600E and ATP concentrations pushes the 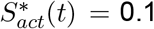 surface towards higher drug doses. In other words, increasing BRAFV600E and ATP concentrations makes DBF-TMT-SCH treatment strategies less effective in silico. This result suggests that high BRAFV600E and ATP concentrations may result in drug resistance to DBF-TMT-SCH vertical inhibition, mediated by intrinsic mechanisms of the BRAFV600E-MEK-ERK pathway. These observations are in line with the discussions that followed the monotherapy simulations, where we examined the impact of BRAFV600E and ATP concentrations on pathway dynamics.

## Discussion

In this study, we present a mathematical model of signalling dynamics in the intracellular BRAFV600E-MEK-ERK pathway. The model can be used to simulate pathway dynamics in response to combinations of the BRAFV00E-inhibitor DBF, the MEK-inhibitor TMT, and the ERK-inhibitor SCH. In its versatility, the model can be used to identify a range of one-, two-, and three-component vertical inhibition treatment strategies that achieve some desired treatment response, such as suppressing the activity of selected pathway components to chosen levels. In this article, we specifically use the model to identify treatment strategies that achieve 90% inhibition of the ERK-activated substrate c-Myc (S in Fig. 4). We argue that this in silico methodology of identifying viable treatment strategies can be used to inform which treatment strategies warrant further investigation in vitro and in vivo. This is important, as it is possible to design more treatment strategies than are feasibly testable in experimental and clinical settings by varying *e*.*g*., drug combinations, drug doses, and drug schedules (although the latter is not discussed in this article). As such, our study contributes an approach to limit the search space for effective vertical inhibition strategies.

**Fig. 4:**
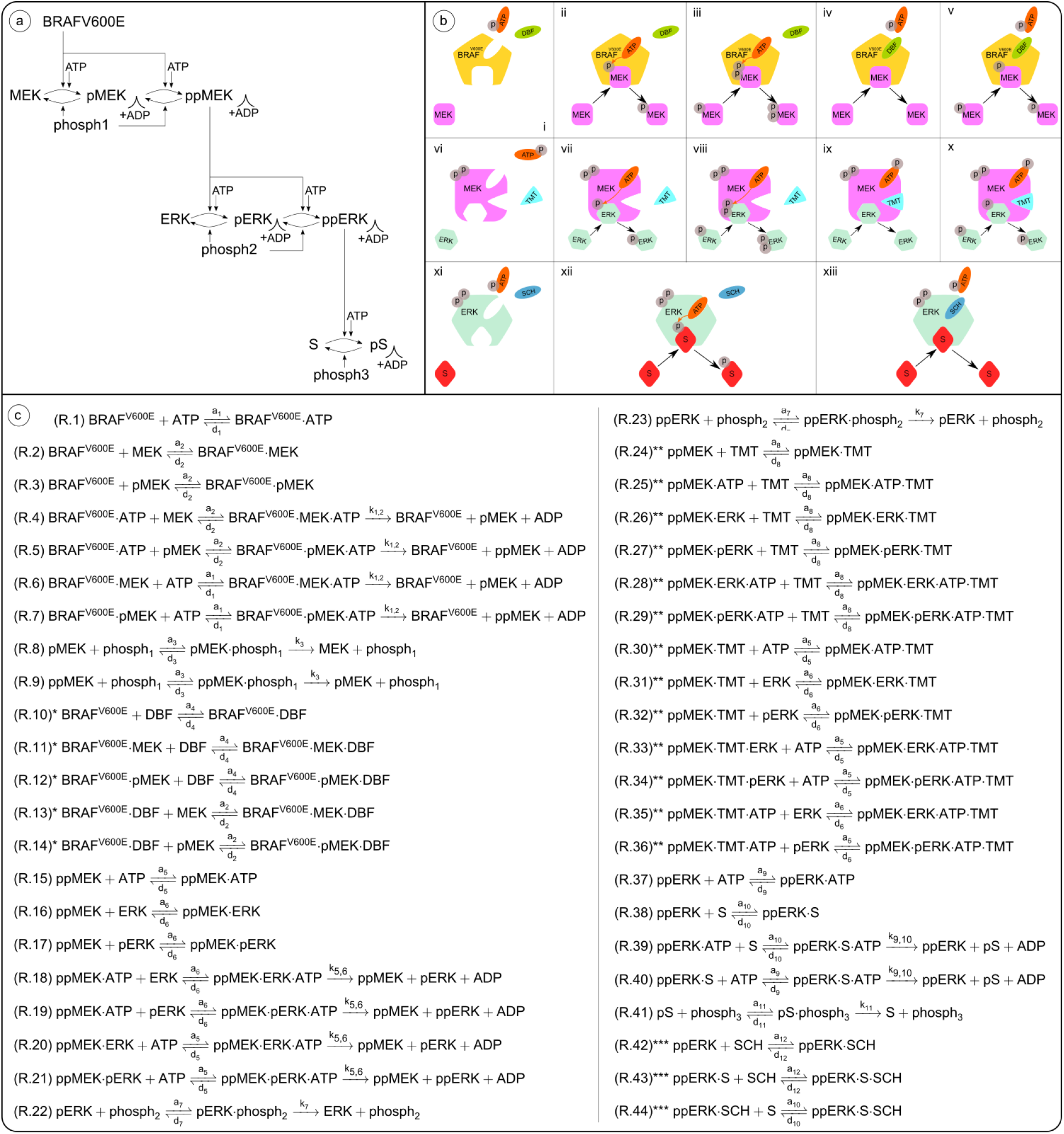
The mathematical model used in this study is composed of a model of the BRAFV600E-MEK-ERK signalling pathway, a set of possible interactions that can occur between pathway components and inhibitors, and a system of chemical reactions that is extracted from the pathway and interaction models. **(a)** The pathway model comprises the BRAFV600E-MEK-ERK signalling cascade and an ERK-activated substrate, S. Singly and doubly phosphorylated pathway components are, respectively, indicated by the prefixes (p) and (pp). Via sequential ATP-meditated phosphorylation events, in which ATP donates a phosphate group to a substrate and turns into ADP, BRAFV600E activates MEK to the form ppMEK. Similarly, ppMEK activates ERK to the form ppERK, which in turn activates the substrate S. Dephosphorylation is mediated by phosphatases (phosph1, phosph2, phosph3). **(b)** The model includes a set of possible interactions that can occur between pathway components and the BRAFV600E-inhibitor DBF (i), the MEK-inhibitor TMT (vi), and the ERK-inhibitor SCH (xi). If inhibitors are not bound to the enzymes, enzyme-bound ATP donates a phosphate group to the substrate MEK (ii)/pMEK (iii)/ERK (vii)/pERK (viii)/S (xii). The ATP-competitive inhibitor DBF binds to the same site as ATP on the BRAFV600E molecule and thus prevents ATP from binding to BRAFV600E and donating a phosphate group to MEK (iv)/pMEK (v). The ATP-allosteric inhibitor TMT binds to a binding site adjacent to the ATP-binding site on the ppMEK molecule and inhibits ATP from donating a phosphate group to ERK (ix)/pERK (x). The ATP-competitive inhibitor SCH binds to the same site as ATP on the ppERK molecule and thus prevents ATP from binding to ppERK and donating a phosphate group to S (xiii). **(c)** A system of chemical reactions is extracted from (a) and (b). Reactions marked by one (*), two (**), or three (***) stars involve the inhibitors DBF, MEK, and SCH, respectively, and can be omitted when modelling pathway dynamics in the absence of drugs. This figure has been adapted from one of our previous publications,^30^ licensed under CC-BY-4.0 (http://creativecommons.org/licenses/by/4.0/)

The methodology outlined in this study can also be used to identify vertical inhibition treatment strategies that comprise two (Fig. 2) or three (Fig. 3) inhibitors that are synergistic in silico. The model identified a broad range of synergistic DBF-TMT dose combinations, but no synergistic DBF-SCH or TMT-SCH doses (Figs. 2 and SM6.1.). However, drug combinations of DBF-TMT,^43^ DBF-SCH,^44^ and DBF-TMT-SCH984^19^ (where SCH984, like SCH772984, is an ATP-competitive ERK-inhibitor) have all shown markedly improved treatment effects, compared to the respective monotherapies alone, in pre-clinical experiments.

The fact that the mathematical model presented in this paper predicts striking DBF-TMT synergies, but no synergies between other inhibitor combinations, suggests that, whilst the pathway model (Fig. 4a) suffices to explain DBF-TMT synergies, a model modification or extension is needed to explain experimentally observed DBF-SCH synergies. Such an extension could, for instance, include drug-protein interactions with un-phosphorylated or singly phosphorylated proteins, so that TMT could bind to MEK, and pMEK (not solely ppMEK), and SCH could bind to ERK and pERK (not solely ppERK). Another possible model extension would be to include alternative pathway components that feed into MEK, or feedback loops in the BRAFV600E-MEK-ERK signalling pathway that are activated by DBF monotherapies but suppressed by combination therapies. On this note, one of the working hypotheses for why vertical pathway inhibition yields better treatment responses than DBF alone is that BRAFV600E-inhibitor monotherapies can activate alternative signalling programs to phosphorylate MEK and ERK.^45^ This type of alternative pathway activation has been recognized as a mechanism of drug resistance to BRAFV600E-inhibitors.^45^ Building on this, Kholodenko et al.^29^ investigated drug resistance mediated by the reactivation of initially inhibited signalling pathways in a recent publication. They analyzed how pathway topologies affect pathway signalling in responses to drug treatments. As an example, they considered RAF-MEK-ERK signalling in BRAFV600E mutated melanoma subjected to vertical inhibition treatment strategies.

Another intracellular mechanism for resistance to BRAFV600E-inhibitors is elevated BRAFV00E activity.^46,47^ This mode of drug resistance is captured by our model, as our simulation results show that increasing BRAFV600E concentrations yield decreasing sensitivity to inhibitors that target the BRAFV600E-MEK-ERK pathway, as illustrated in Figs. 1 and 3. These simulation results also show that ATP concentrations impact drug resistance in silico, although we note that the clinical relevance of this finding is not straightforward to interpret. This is because tumour cells in physiological oxygen conditions decrease their oxidative phosphorylation in favour of increased use of glycolysis, according to the Warburg effect.^48,49,50^ Though both processes produce ATP, which generates energy for cellular processes, oxidative phosphorylation generally generates ATP more effectively than glycolysis.^51^ The complex relationship between drug resistance, ATP, and ATP production in tumour cells is beyond the scope of this study but has been investigated in other works.^51,52,53^ We also remark that the discrepancy in ATP levels between experimental models and in situ tumours has been suggested to contribute to the discrepancy in drug performance between laboratory and physiological settings.^52^ Our results, which show that ATP levels may impact treatment responses to ATP-competitive and ATP-allosteric inhibitors in silico, support this argument.

ERK pathways have been studied with a variety of mathematical models of different spatial and temporal resolutions. This is, in large part, due to their importance for normal cell functions, and their links to diseases such as cancer, developmental disorders and neurological disorders.^11^ In the “Introduction” section of this article, we discussed a handful of mathematical ERK models that are directly related to the design of our study. Here, we describe a selection of other ERK and BRAFV600E-related models that demonstrates a range of mathematical, statistical and computation techniques. Gerosa et al. combined mathematical modelling with cell imaging and proteomics to study how BRAFV600E-mutated melanoma cells adapt to drugs. They showed that drugs induce pathway re-wiring which results in sporadic ERK pulses that promote cell survival and proliferation.^54^ Pillai et al. demonstrated the existence of phenotypic multi-stability in melanoma with a gene regulatory network model. The model was related to drug resistance and provided explanations for sub-cellular mechanisms that drive phenotypic heterogeneity and phenotypic switching in melanoma.^55^ Smalley et al. showed that information about cell-level transcriptional heterogeneity in melanoma xenografts can be used to predict initial treatment responses to BRAF-inhibitors. They used a two-compartment ODE model, in which the two compartments represented the amounts of drug-sensitive and drug-resistant cells in the xenografts.^56^ Lai and Friedman used a partial differential equation model that described the dynamics of cancer cells, T cells, proinflammatory cytokines, a BRAF/MEK inhibitor, and a drug that targets the immunoinhibitory receptor PD-1 on T cells to show that combination treatments with the two drugs are positively correlated at low doses, but are antagonistic for some high doses in silico.^57^ Sun et al. studied therapy-induced drug resistance in melanoma with a stochastic differential equation model. The model revealed dose-dependent synergy of combination treatments that involve BRAF, MEK, and PI3K inhibitors, and suggested that optimal dose combination therapies may reduce drug resistance.^58^ Marsh et al. used model reduction techniques, including those based on algebraic topology, to reduce an ERK reaction network model. Such methods are useful as mathematical models of intracellular pathway dynamics can become large, which may complicate model analysis and parameter estimation.^59^ Moreover, the experimental and computational modelling studies by Schoeberl et al.^28^ and Fujioka et al.^60^ have made important contributions to the estimation and reporting of parameter values that pertain to MAPK-cascade related rate constants and kinase concentrations in human cells.

On that note, we remark that one limiting factor of this study is that the model parameter values are gathered and derived from multiple literature sources (Supplementary Material, Supplementary Note 3). As such, the parameter values are based on results from different studies with different experimental settings, which we need to consider when evaluating the values’ correctness. We anticipate that access to a complete set of cell-line-specific model parameter values, derived from one baseline experimental setting, would improve the predictive ability of the model. Just as importantly, such a setting would (a) provide an experimental system in which to test and evaluate the model’s predictive performance, and (b) inform model updates. In our previous study, we evaluated model-predicted levels of activated ERK in response to DBF monotherapies against data from clinical samples (Fig. 6 in Hamis et al., 2021^30^). The results of this evaluation were promising, which indicates that the model used in our previous study, and by extension the model used in our current study, has the potential to be of clinical relevance. Here, our model is designed to focus on signalling dynamics that result from interactions between components of the BRAFV600E-MEK-ERK pathway and three drugs: DBF, TMT and SCH. As such, our model simulates intracellular signalling dynamics within one cell as a closed system. To more realistically represent in vivo or clinical signalling dynamics, the model could be extended to include *e*.*g*., drug transportation, drug elimination, immune responses, dynamic drug resistance and cell crowding effects. Challenges and techniques for bridging mathematical in vitro findings with mouse-model and clinical research have been discussed in previous bio-mathematical articles.^61,62^ Even in its current form, our model is able to qualitatively capture important aspects of vertical inhibition of the BRAFV600E-MEK-ERK pathway that have been observed in vitro. Notably, the model shows that DBF-TMT combination therapies may synergistically inhibit ERK-activated substrate phosphorylation and that SCH therapies are effective in both the presence and absence of inhibitors that target BRAFV600E and MEK.^36,19^

On the topic of translational and clinical relevance, we remark that beyond BRAFV600E-mutant melanoma, the BRAFV600E-MEK-ERK pathway is also a target for treating non-small-cell lung carcinoma, thyroid cancer, and paediatric high-grade glioma with the BRAFV600E-mutation.^63,64^ The mathematical model presented in this article could be modified, ideally by cancer-type specific model parameter values, to simulate BRAFV600E-MEK-ERK pathway dynamics and vertical inhibition treatment responses in other cancers. The model could also be altered to study the dynamics of other ERK-activated substrates such as the transcription factors c-Fos and Elk1, which also impact carcinogenesis.^11^

To summarise, the translational value of this bio-mathematical study is two-fold. Firstly, the mathematical model increases our understanding of BRAFV600E-MEK-ERK signalling dynamics in response to vertical inhibition. Notably, the model provides data-driven mechanistic modelling explanations for why resistance to DBF monotherapies and DBF-TMT-SCH combination therapies markedly increases with intracellular BRAFV600E concentrations, whilst resistance to TMT and SCH monotherapies does not. Secondly, the methodology presented in this study can be used as a tool to inform the design of pre-clinical studies that aim to find a range of drug dose combinations that target the BRAFV600E-MEK-ERK signalling pathway and achieve desired treatment responses. These mathematically identified dose combinations could offer alternatives to high-dose combinations which are currently used in clinics. As such, this bio-mathematical study constitutes a calculated step towards rational treatment strategies.

## Methods

The mathematical model used in this study is composed of a pathway model of the intracellular BRAF V600E-MEK-ERK pathway with the substrate c-Myc (S) downstream of ERK (Fig 4a), and a set of possible interactions that can occur between pathway components and inhibitors (Fig 4b). When ERK phosphorylates c-Myc at the Ser62 phosphorylation site, c-Myc expression is stabilized.^35^ In melanoma, stable c-Myc overexpression has been observed to promote cell proliferation and is associated with poor prognosis.^65^ Accordingly, we choose phosphorylated c-Myc (pS in Fig. 4) to be our main model output of interest, and the simulated treatments ultimately act to suppress the amount of pS in the modelled system. From the pathway model and the set of possible interactions, we extract a system of chemical reactions that describe pathway signalling in response to vertical inhibition (Fig 4c). Using a computational framework^30^ that builds on the law of mass action, we convert the system of chemical reactions into a system of ODEs that are solved numerically to produce simulations of pathway signalling dynamics. The model is adapted from previous studies,^25,30^ and is extended to include an ERK-activated substrate S (c-Myc) and an ERK-inhibitor (SCH772984). The next subsections provide detailed descriptions of the mathematical model and the computational methods used in this study. Information on how to access, run, and modify the code files is available in the Supplementary Material (Supplementary Note 2).

### Modelling the BRAFV600E-MEK-ERK pathway in the absence of inhibitors

The pathway model consists of a three-tiered cascade comprising BRAFV600E, and various forms of MEK, ERK, and the ERK-activated substrate S (Fig 4a). Throughout this study, single (p) and double (pp) p-prefixes respectively indicate that one or two phosphate groups are bound to the pathway component of interest. At the top of the cascade is the BRAFV600E-mutated BRAF oncogene, which we model as always activated to simulate hyper-activated pathway signalling. BRAFV600E can transform MEK into its activated form ppMEK, which can transform ERK into its activated form ppERK, which in turn can activate the substrate S to the form pS. More precisely, activation events downstream of BRAFV600E are mediated via one or two sequential phosphorylation events, where an ATP molecule binds to an enzyme (*i*.*e*. BRAFV600E/ppMEK/ppERK) and donates a phosphate group to a substrate (*i*.*e*. MEK/pMEK/ERK/pERK/S) (Fig 4a,b). After the phosphate group donation, the ATP molecule converts to adenosine diphosphate (ADP). In our model, dephosphorylation is mediated by the phosphatases phosph1, phosph2, and phosph3 which, respectively, remove phosphate groups from the molecules pMEK/ppMEK, pERK/ppERK, and pS (Fig 4a,c).

The pathway model used in this study is an extension of our recently published model of the BRAFV600E-MEK-ERK patwhay,^30^ which is an adaptation of the more general pathway model presented by Huang and Ferrell in 1996.^25^ The components downstream of ppERK in the pathway model are introduced in the current study (Fig. 4a).

### Modelling interactions between pathway components and inhibitors

We model interactions between pathway components and inhibitors on a molecular level (Fig 4b). To do this, we distinguish between ATP-competitive and ATP-allosteric inhibitors. DBF^43^ and SCH^36^ are ATP-competitive inhibitors, meaning that they compete with ATP for a binding site on their target enzyme, *i*.*e*. BRAFV600E, and ppERK, respectively. As such, DBF prevents ATP from binding to BRAFV600E and, by extension, inhibits ATP-mediated MEK phosphorylation. Similarly, SCH prevents ATP from binding to ppERK and, by extension, inhibits ATP-mediated S phosphorylation. On the other hand, TMT is an ATP-allosteric inhibitor,^66^ and thus TMT binds to a MEK binding site that is adjacent to, but distinct from, the ATP-specific binding site. MEK-bound TMT prevents MEK-bound ATP from donating a phosphate group to MEK-bound ERK or pERK. The modelled interactions pertaining to BRAFV600E and MEK inhibition (Fig. 4b, i-x) were presented in one of our previous studies, whereas the interactions pertaining to ERK inhibition are introduced in this study (Fig. 4b, xi-xiii). Note that we have made the simplifying modelling assumption that the drugs TMT and SCH only bind to activated, *i*.*e*., doubly phosphorylated, forms of MEK and ERK, respectively.

### Formulating the system of chemical reactions that describes pathway dynamics

From the pathway model (Fig. 4a) and the set of possible interactions between pathway components and inhibitors (Fig. 4b), we extract a system of chemical reactions (Fig. 4c). These reactions are associated with rate constants that we obtain from published in vitro data, as described in the Supplementary Material (Supplementary Note 3).

### Converting chemical reactions to equations, and simulating pathway dynamics

Using the law of mass action, we convert the system of chemical reactions R.1-R.44 (Fig. 4) to a system of ODEs. The law of mass action states that the rate of a chemical reaction is proportional to the product of the reactants’ concentrations.^67^ The reactions-to-ODEs conversion process is automated by a computational MATLAB^68^ framework, that we (some of the authors) previously developed.^30^ The full system of ODEs for our model (Fig. 4) is available in the Supplementary Material (Supplementary Note 1.2). After formulating the equations, we also use the computational framework to numerically solve the equations and thus produce simulations of intracellular pathway dynamics. The simulations show how concentrations of the pathway components change over time. In this study, we are interested in studying vertical inhibition strategies in which up to three components of the BRAFV600E-MEK-ERK signalling pathway are simultaneously inhibited. Therefore, a specific output of interest is the fraction of activated (*i*.*e*., simultaneously free and phosphorylated) substrate S in the system which we call *S*_*act*_ and define as

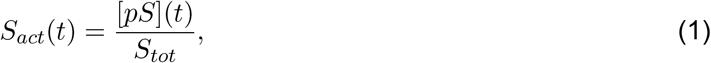

where we have used the slanted bracket notation [*x*](*t*) to denote the concentration of substrate *x* at time *t*, and *S*_*tot*_ is the total amount of substrate S in the system. Our mathematical model obeys a set of conservation laws that ensures the total amount of various pathway components is constant throughout the simulations. For example, the conservation law for *S*_*tot*_ reads

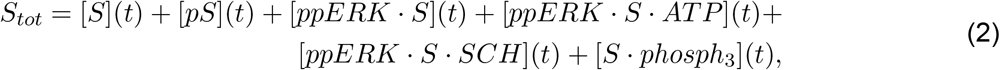

where the left-hand-side of Eq. 2 corresponds to the denominator in Eq. 1. The full list of conservation laws is available in the Supplementary Material (Supplementary Note 1.3). Building on Eq. 1, we introduce the variable 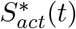 to clearly quantify the inhibitory effects of in silico treatments. We set

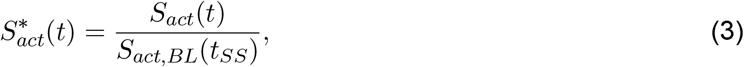

where *S*_*act,BL*_(*t*) denotes *S*_*act*_(*t*) computed for baseline (*BL*) intracelullar BRAFV600E and ATP concentrations, and in the absence of drugs. Moreover, *t*_*SS*_ denotes the time it takes for *S*_*act,BL*_(*t*) to reach a steady-state. In this study, we thus say that a treatment strategy that yields 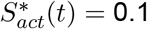 achieves a 90% inhibition of substrate S activity.

### Using in vitro data to parameterize the model

The system of reactions (Fig. 4c, R.1-R.44) includes 12 forward rate constants *a*_1_, *a*_2_, …, *a*_12_, 12 reverse rate constants *d*_1_, *d*_2_, …, *d*_12_ and 6 catalytic rate constants *k*_1,2_, *k*_3_, *k*_5,6_, *k*_7_, *k*_9,10_, *k*_11_. The rate constants are listed in the Supplementary Material (Supplementary Table 1) and are derived from published data^25,35,36,69,70,71,72,73,74,75,76,77^ using binding data calculations^78,79,80^ (SM3.1,3). The initial and total concentrations of all pathway components are listed in the Supplementary Material (Supplementary Table 2) and are obtained from published data^25,75^ (Supplementary Note 3.2).

## Data availability

All simulation data are available on the code-hosting platform GitHub (https://github.com/SJHamis/MAPKcascades). Instructions on how to access, run, and modify the code files are available in the Supplementary Material (Supplementary Note 2).

## Code availability

All code files are available on the code-hosting platform GitHub (https://github.com/SJHamis/MAPKcascades). Instructions on how to access, run, and modify the code files are available in the Supplementary Material (Supplementary Note 2).

## Acknowledgements

SH was financially supported by Wenner-Gren Stiftelserna/the Wenner-Gren Foundations (WGF2022-0044), the Tampere Institute for Advanced Study (2021-2023), and the Kjell och Märta Beijer Foundation.

## Authors contributions

ADC and SH produced the results and drafted the manuscript. SH supervised modelling, coding and manuscript-drafting. All authors contributed to the design of the work. All authors revised and have approved of the manuscript, and have agreed to be accountable for all aspects of the work presented in the manuscript.

## Competing interests

The authors have no competing interests to declare that are relevant to the content of this article.

## Notes

https://github.com/SJHamis/MAPKcascades

### Competing Interest Statement

The authors have declared no competing interest.

### Summary of Updates

The order of the sections has been updated. The labeling of Supplementary Material content has been update. Figure 1 has been polished.

