## Supplementary Notes 1-6 for "Simulating BRAFV600E-MEK-ERK signalling dynamics in response to vertical inhibition treatment strategies"

3.3.3 Estimating rate constants for substrate-phosphatase interactions 12
3.3.4 Estimating rate constants for ppERK-SCH interactions . . . . . 12

**4 Supplementary Note 4: Demonstrating the transient behaviour of system dynamics in response to SCH monotherapies** **13**

**5 Supplementary Note 5: Identifying two-drug combinations that yield synergistic responses** **13**

**6 Supplementary Note 6: Identifying three-component vertical inhibition strategies that achieve desired substrate activation levels** **14**

**Supplementary References** **19**

### 1 Supplementary Note 1: Formulating the system of differential-algebraic equations

#### 1.1 The system of reactions (R.1-R.44)

The system of chemical reactions (R.1-R.44) describes signalling dynamics in the intracellular BRAFV600E-MEK-ERK pathway in response to the BRAFV600E-inhibitor dabrafenib (DBF), the MEK-inhibitor trametinib (TMT) and the ERK-inhibitor SCH772984 (SCH). The pathway and drug actions are schematically illustrated in Fig.1 in the main manuscript. The dot notation ($\cdot$) is used to denote complexes of two or more molecules. Reactions involving the BRAFV600E, MEK, and ERK inhibitors are respectively marked by single (*), double (**), and triple (***) stars, and can be omitted to study the system in the absence of drugs. Note that reactions (R.1-R.36) were first presented in our previous article,[1] whilst reactions (R.37-R.44) are introduced in the current article.

**System of reactions** (R.1-R.44):

(R.1) $\mathrm{BRAF^{V600E} + ATP \underset{d_1}{\overset{a_1}{\rightleftharpoons}} BRAF^{V600E}{\cdot}ATP}$

(R.2) $\mathrm{BRAF^{V600E} + MEK \underset{d_2}{\overset{a_2}{\rightleftharpoons}} BRAF^{V600E}{\cdot}MEK}$

(R.3) $\mathrm{BRAF^{V600E} + pMEK \underset{d_2}{\overset{a_2}{\rightleftharpoons}} BRAF^{V600E}{\cdot}pMEK}$

(R.4) $\mathrm{BRAF^{V600E}{\cdot}ATP + MEK \underset{d_2}{\overset{a_2}{\rightleftharpoons}} BRAF^{V600E}{\cdot}MEK{\cdot}ATP \xrightarrow{k_{1,2}} BRAF^{V600E} + pMEK + ADP}$

(R.5) $\mathrm{BRAF^{V600E}{\cdot}ATP + pMEK \underset{d_2}{\overset{a_2}{\rightleftharpoons}} BRAF^{V600E}{\cdot}pMEK{\cdot}ATP \xrightarrow{k_{1,2}} BRAF^{V600E} + ppMEK + ADP}$

(R.6) $\mathrm{BRAF^{V600E}{\cdot}MEK + ATP \underset{d_1}{\overset{a_1}{\rightleftharpoons}} BRAF^{V600E}{\cdot}MEK{\cdot}ATP \xrightarrow{k_{1,2}} BRAF^{V600E} + pMEK + ADP}$

(R.7) $\mathrm{BRAF^{V600E}{\cdot}pMEK + ATP \underset{d_1}{\overset{a_1}{\rightleftharpoons}} BRAF^{V600E}{\cdot}pMEK{\cdot}ATP \xrightarrow{k_{1,2}} BRAF^{V600E} + ppMEK + ADP}$

(R.8) $\mathrm{pMEK + phosph_1 \underset{d_3}{\overset{a_3}{\rightleftharpoons}} pMEK{\cdot}phosph_1 \xrightarrow{k_3} MEK + phosph_1}$

(R.9) $\mathrm{ppMEK + phosph_1 \underset{d_3}{\overset{a_3}{\rightleftharpoons}} ppMEK{\cdot}phosph_1 \xrightarrow{k_3} pMEK + phosph_1}$

(R.10)* $\mathrm{BRAF^{V600E} + DBF \underset{d_4}{\overset{a_4}{\rightleftharpoons}} BRAF^{V600E}{\cdot}DBF}$

(R.11)* $\mathrm{BRAF^{V600E}{\cdot}MEK + DBF \underset{d_4}{\overset{a_4}{\rightleftharpoons}} BRAF^{V600E}{\cdot}MEK{\cdot}DBF}$

(R.12)* $\mathrm{BRAF^{V600E}{\cdot}pMEK + DBF \underset{d_4}{\overset{a_4}{\rightleftharpoons}} BRAF^{V600E}{\cdot}pMEK{\cdot}DBF}$

(R.13)* $\mathrm{BRAF^{V600E}{\cdot}DBF + MEK \underset{d_2}{\overset{a_2}{\rightleftharpoons}} BRAF^{V600E}{\cdot}MEK{\cdot}DBF}$

(R.14)* $\mathrm{BRAF^{V600E}{\cdot}DBF + pMEK \underset{d_2}{\overset{a_2}{\rightleftharpoons}} BRAF^{V600E}{\cdot}pMEK{\cdot}DBF}$

(R.15) $\mathrm{ppMEK + ATP \underset{d_5}{\overset{a_5}{\rightleftharpoons}} ppMEK{\cdot}ATP}$

(R.16) $\mathrm{ppMEK + ERK \underset{d_6}{\overset{a_6}{\rightleftharpoons}} ppMEK{\cdot}ERK}$

(R.17) $\mathrm{ppMEK + pERK \underset{d_6}{\overset{a_6}{\rightleftharpoons}} ppMEK{\cdot}pERK}$

(R.18) $\mathrm{ppMEK{\cdot}ATP + ERK \underset{d_6}{\overset{a_6}{\rightleftharpoons}} ppMEK{\cdot}ERK{\cdot}ATP \xrightarrow{k_{5,6}} ppMEK + pERK + ADP}$

(R.19) $\mathrm{ppMEK{\cdot}ATP + pERK \underset{d_6}{\overset{a_6}{\rightleftharpoons}} ppMEK{\cdot}pERK{\cdot}ATP \xrightarrow{k_{5,6}} ppMEK + ppERK + ADP}$

(R.20) $\mathrm{ppMEK{\cdot}ERK + ATP \underset{d_5}{\overset{a_5}{\rightleftharpoons}} ppMEK{\cdot}ERK{\cdot}ATP \xrightarrow{k_{5,6}} ppMEK + pERK + ADP}$

(R.21) $\mathrm{ppMEK{\cdot}pERK + ATP \underset{d_5}{\overset{a_5}{\rightleftharpoons}} ppMEK{\cdot}pERK{\cdot}ATP \xrightarrow{k_{5,6}} ppMEK + ppERK + ADP}$

(R.22) $\mathrm{pERK + phosph_2 \underset{d_7}{\overset{a_7}{\rightleftharpoons}} pERK{\cdot}phosph_2 \xrightarrow{k_7} ERK + phosph_2}$

(R.23) ppERK + $phosph_2 \underset{d_7}{\overset{a_7}{\rightleftharpoons}}$ ppERK·$phosph_2 \xrightarrow{k_7}$ pERK + $phosph_2$

(R.24)** ppMEK + TMT $\underset{d_8}{\overset{a_8}{\rightleftharpoons}}$ ppMEK·TMT

(R.25)** ppMEK·ATP + TMT $\underset{d_8}{\overset{a_8}{\rightleftharpoons}}$ ppMEK·ATP·TMT

(R.26)** ppMEK·ERK + TMT $\underset{d_8}{\overset{a_8}{\rightleftharpoons}}$ ppMEK·ERK·TMT

(R.27)** ppMEK·pERK + TMT $\underset{d_8}{\overset{a_8}{\rightleftharpoons}}$ ppMEK·pERK·TMT

(R.28)** ppMEK·ERK·ATP + TMT $\underset{d_8}{\overset{a_8}{\rightleftharpoons}}$ ppMEK·ERK·ATP·TMT

(R.29)** ppMEK·pERK·ATP + TMT $\underset{d_8}{\overset{a_8}{\rightleftharpoons}}$ ppMEK·pERK·ATP·TMT

(R.30)** ppMEK·TMT + ATP $\underset{d_5}{\overset{a_5}{\rightleftharpoons}}$ ppMEK·ATP·TMT

(R.31)** ppMEK·TMT + ERK $\underset{d_6}{\overset{a_6}{\rightleftharpoons}}$ ppMEK·ERK·TMT

(R.32)** ppMEK·TMT + pERK $\underset{d_6}{\overset{a_6}{\rightleftharpoons}}$ ppMEK·pERK·TMT

(R.33)** ppMEK·TMT·ERK + ATP $\underset{d_5}{\overset{a_5}{\rightleftharpoons}}$ ppMEK·ERK·ATP·TMT

(R.34)** ppMEK·TMT·pERK + ATP $\underset{d_5}{\overset{a_5}{\rightleftharpoons}}$ ppMEK·pERK·ATP·TMT

(R.35)** ppMEK·TMT·ATP + ERK $\underset{d_6}{\overset{a_6}{\rightleftharpoons}}$ ppMEK·ERK·ATP·TMT

(R.36)** ppMEK·TMT·ATP + pERK $\underset{d_6}{\overset{a_6}{\rightleftharpoons}}$ ppMEK·pERK·ATP·TMT

(R.37) ppERK + ATP $\underset{d_9}{\overset{a_9}{\rightleftharpoons}}$ ppERK·ATP

(R.38) ppERK + S $\underset{d_{10}}{\overset{a_{10}}{\rightleftharpoons}}$ ppERK·S

(R.39) ppERK·ATP + S $\underset{d_{10}}{\overset{a_{10}}{\rightleftharpoons}}$ ppERK·S·ATP $\xrightarrow{k_{9,10}}$ ppERK + pS + ADP

(R.40) ppERK·S + ATP $\underset{d_9}{\overset{a_9}{\rightleftharpoons}}$ ppERK·S·ATP $\xrightarrow{k_{9,10}}$ ppERK + pS + ADP

(R.41) pS + $phosph_3 \underset{d_{11}}{\overset{a_{11}}{\rightleftharpoons}}$ pS·$phosph_3 \xrightarrow{k_{11}}$ S + $phosph_3$

(R.42)*** ppERK + SCH $\underset{d_{12}}{\overset{a_{12}}{\rightleftharpoons}}$ ppERK·SCH

(R.43)*** ppERK·S + SCH $\underset{d_{12}}{\overset{a_{12}}{\rightleftharpoons}}$ ppERK·S·SCH

(R.44)*** ppERK·SCH + S $\underset{d_{10}}{\overset{a_{10}}{\rightleftharpoons}}$ ppERK·S·SCH

#### 1.2 The system of ordinary differential equations (O.1-O.46)

Using the law of mass action, the system of reactions (R.1-R.44) is converted to a system of ordinary differential equations (O.1-O.46). This reactions-to-ODEs conversion is automated by a computational framework that we (some of the authors) previously presented.[1] This framework is discussed in Section 2.

To enable straight-forward implementation of the ODE system, we let $\mathbf{y_{46\times1}}$ denote a column vector with 46 elements, where each element corresponds to a signalling molecule concentration so that,

$$
\begin{aligned}
y(1) &= [BRAFV600E],\\
y(2) &= [ATP],\\
y(3) &= [ATP \cdot BRAFV600E],\\
y(4) &= [MEK],\\
y(5) &= [BRAFV600E \cdot MEK],\\
y(6) &= [pMEK],
\end{aligned}
$$

$$
\begin{aligned}
y(7) &= [BRAFV600E \cdot pMEK], \\
y(8) &= [ATP \cdot BRAFV600E \cdot MEK], \\
y(9) &= [ADP], \\
y(10) &= [ATP \cdot BRAFV600E \cdot pMEK], \\
y(11) &= [ppMEK], \\
y(12) &= [phosph_1], \\
y(13) &= [pMEK \cdot phosph_1], \\
y(14) &= [phosph_1 \cdot ppMEK], \\
y(15) &= [DBF], \\
y(16) &= [BRAFV600E \cdot DBF], \\
y(17) &= [BRAFV600E \cdot DBF \cdot MEK], \\
y(18) &= [BRAFV600E \cdot DBF \cdot pMEK], \\
y(19) &= [ATP \cdot ppMEK], \\
y(20) &= [ERK], \\
y(21) &= [ERK \cdot ppMEK], \\
y(22) &= [pERK], \\
y(23) &= [pERK \cdot ppMEK], \\
y(24) &= [ATP \cdot ERK \cdot ppMEK], \\
y(25) &= [ATP \cdot pERK \cdot ppMEK], \\
y(26) &= [ppERK], \\
y(27) &= [phosph_2], \\
y(28) &= [pERK \cdot phosph_2], \\
y(29) &= [phosph_2 \cdot ppERK], \\
y(30) &= [TMT], \\
y(31) &= [TMT \cdot ppMEK], \\
y(32) &= [ATP \cdot TMT \cdot ppMEK], \\
y(33) &= [ERK \cdot TMT \cdot ppMEK], \\
y(34) &= [TMT \cdot pERK \cdot ppMEK], \\
y(35) &= [ATP \cdot ERK \cdot TMT \cdot ppMEK], \\
y(36) &= [ATP \cdot TMT \cdot pERK \cdot ppMEK], \\
y(37) &= [ATP \cdot ppERK], \\
y(38) &= [S], \\
y(39) &= [S \cdot ppERK], \\
y(40) &= [ATP \cdot S \cdot ppERK], \\
y(41) &= [pS], \\
y(42) &= [phosph_3], \\
y(43) &= [pS \cdot phosph_3], \\
y(44) &= [SCH], \\
y(45) &= [SCH \cdot ppERK], \\
y(46) &= [SCH \cdot S \cdot ppERK].
\end{aligned}
$$

Using this notation, the system of ODEs (O.1-O.46) can be formulated as,

$$\frac{dy(1)}{dt} = y(1)y(2)(-a_1) + d_1y(3) + y(1)y(4)(-a_2) + d_2y(5) + y(1)y(6)(-a_2) + d_2y(7) + y(8)k_{12} + y(10)k_{12} + y(8)k_{12} + y(10)k_{12} + y(1)y(15)(-a_4) + d_4y(16), \tag{O.1}$$

$$\frac{dy(2)}{dt} = y(1)y(2)(-a_1) + d_1y(3) + y(5)y(2)(-a_1) + d_1y(8) + y(7)y(2)(-a_1) + d_1y(10) + y(11)y(2)(-a_5) + d_5y(19) + y(21)y(2)(-a_5) + d_5y(24) + y(23)y(2)(-a_5) + d_5y(25) + y(31)y(2)(-a_5) + d_5y(32) + y(33)y(2)(-a_5) + d_5y(35) + y(34)y(2)(-a_5) + d_5y(36) + y(26)y(2)(-a_9) + d_9y(37) + y(39)y(2)(-a_9) + d_9y(40), \tag{O.2}$$

$$\frac{dy(3)}{dt} = y(3)(-d_1) + a_1(y(1)y(2)) + y(3)y(4)(-a_2) + d_2y(8) + y(3)y(6)(-a_2) + d_2y(10), \tag{O.3}$$

$$\frac{dy(4)}{dt} = y(1)y(4)(-a_2) + d_2y(5) + y(3)y(4)(-a_2) + d_2y(8) + y(13)k_3 + y(16)y(4)(-a_2) + d_2y(17), \tag{O.4}$$

$$\frac{dy(5)}{dt} = y(5)(-d_2) + a_2(y(1)y(4)) + y(5)y(2)(-a_1) + d_1y(8) + y(5)y(15)(-a_4) + d_4y(17), \tag{O.5}$$

$$\frac{dy(6)}{dt} = y(1)y(6)(-a_2) + d_2y(7) + y(8)k_{12} + y(3)y(6)(-a_2) + d_2y(10) + y(8)k_{12} + y(6)y(12)(-a_3) + d_3y(13) + y(14)k_3 + y(16)y(6)(-a_2) + d_2y(18), \tag{O.6}$$

$$\frac{dy(7)}{dt} = y(7)(-d_2) + a_2(y(1)y(6)) + y(7)y(2)(-a_1) + d_1y(10) + y(7)y(15)(-a_4) + d_4y(18), \tag{O.7}$$

$$\frac{dy(8)}{dt} = y(8)(-d_2 - k_{12}) + a_2(y(3)y(4)) + y(8)(-d_1 - k_{12}) + a_1(y(5)y(2)) \tag{O.8}$$

$$\frac{dy(9)}{dt} = y(8)k_{12} + y(10)k_{12} + y(8)k_{12} + y(10)k_{12} + y(24)k_{56} + y(25)k_{56} + y(24)k_{56}k_{56} + y(25)k_{56} + y(40)k_{910} + y(40)k_{910}, \tag{O.9}$$

$$\frac{dy(10)}{dt} = y(10)(-d_2 - k_{12}) + a_2(y(3)y(6)) + y(10)(-d_1 - k_{12}) + a_1(y(7)y(2)), \tag{O.10}$$

$$\frac{dy(11)}{dt} = y(10)k_{12} + y(10)k_{12} + y(11)y(12)(-a_3) + d_3y(14) + y(11)y(2)(-a_5) + d_5y(19) + y(11)y(20)(-a_6) + d_6y(21) + y(11)y(22)(-a_6) + d_6y(23) + y(24)k_{56} + y(25)k_{56} + y(24)k_{56} + y(25)k_{56} + y(11)y(30)(-a_8) + d_8y(31), \tag{O.11}$$

$$\frac{dy(12)}{dt} = y(6)y(12)(-a_3) + d_3y(13) + y(13)k_3 + y(11)y(12)(-a_3) + d_3y(14) + y(14)k_3, \tag{O.12}$$

$$\frac{dy(13)}{dt} = y(13)(-d_3 - k_3) + a_3(y(6)y(12)), \tag{O.13}$$

$$\frac{dy(14)}{dt} = y(14)(-d_3 - k_3) + a_3(y(11)y(12)), \tag{O.14}$$

$$\frac{dy(15)}{dt} = y(1)y(15)(-a_4) + d_4y(16) + y(5)y(15)(-a_4) + d_4y(17) + y(7)y(15)(-a_4) + d_4y(18), \tag{O.15}$$

$$\frac{dy(16)}{dt} = y(16)(-d_4) + a_4(y(1)y(15)) + y(16)y(4)(-a_2) + d_2y(17) + y(16)y(6)(-a_2) + d_2y(18), \tag{O.16}$$

$$\frac{dy(17)}{dt} = y(17)(-d_4) + a_4(y(5)y(15)) + y(17)(-d_2) + a_2(y(16)y(4)), \tag{O.17}$$

$$\frac{dy(18)}{dt} = y(18)(-d_4) + a_4(y(7)y(15)) + y(18)(-d_2) + a_2(y(16)y(6)), \tag{O.18}$$

$$\begin{aligned}\frac{dy(19)}{dt} = {} & y(19)(-d_5) + a_5(y(11)y(2)) + y(19)y(20)(-a_6) + d_6y(24) \\ & + y(19)y(22)(-a_6) + d_6y(25) + y(19)y(30)(-a_8) + d_8y(32),\end{aligned} \tag{O.19}$$

$$\begin{aligned}\frac{dy(20)}{dt} = {} & y(11)y(20)(-a_6) + d_6y(21) + y(19)y(20)(-a_6) + d_6y(24) + y(28)k_7 \\ & + y(31)y(20)(-a_6) + d_6y(33) + y(32)y(20)(-a_6) + d_6y(35),\end{aligned} \tag{O.20}$$

$$\begin{aligned}\frac{dy(21)}{dt} = {} & y(21)(-d_6) + a_6(y(11)y(20)) + y(21)y(2)(-a_5) + d_5y(24) \\ & + y(21)y(30)(-a_8) + d_8y(33),\end{aligned} \tag{O.21}$$

$$\begin{aligned}\frac{dy(22)}{dt} = {} & y(11)y(22)(-a_6) + d_6y(23) + y(24)k_{56} + y(19)y(22)(-a_6) + d_6y(25) \\ & + y(24)k_{56} + y(22)y(27)(-a_7) + d_7y(28) + y(29)k_7 + y(31)y(22)(-a_6) \\ & + d_6y(34) + y(32)y(22)(-a_6) + d_6y(36),\end{aligned} \tag{O.22}$$

$$\begin{aligned}\frac{dy(23)}{dt} = {} & y(23)(-d_6) + a_6(y(11)y(22)) + y(23)y(2)(-a_5) + d_5y(25) \\ & + y(23)y(30)(-a_8) + d_8y(34),\end{aligned} \tag{O.23}$$

$$\begin{aligned}\frac{dy(24)}{dt} = {} & y(24)(-d_6 - k_{56}) + a_6(y(19)y(20)) + y(24)(-d_5 - k_{56}) \\ & + a_5(y(21)y(2)) + y(24)y(30)(-a_8) + d_8y(35),\end{aligned} \tag{O.24}$$

$$\begin{aligned}\frac{dy(25)}{dt} = {} & y(25)(-d_6 - k_{56}) + a_6(y(19)y(22)) + y(25)(-d_5 - k_{56}) \\ & + a_5(y(23)y(2)) + y(25)y(30)(-a_8) + d_8y(36),\end{aligned} \tag{O.25}$$

$$\begin{aligned}\frac{dy(26)}{dt} = {} & y(25)k_{56} + y(25)k_{56} + y(26)y(27)(-a_7) + d_7y(29) + y(26)y(2)(-a_9) \\ & + d_9y(37) + y(26)y(38)(-a_{10}) + d_{10}y(39) + y(40)k_{910} \\ & + y(40)k_{910} + y(26)y(44)(-a_{12}) + d_{12}y(45),\end{aligned} \tag{O.26}$$

$$\begin{aligned}\frac{dy(27)}{dt} = {} & y(22)y(27)(-a_7) + d_7y(28) + y(28)k_7 + y(26)y(27)(-a_7) + d_7y(29) \\ & + y(29)k_7,\end{aligned} \tag{O.27}$$

$$\frac{dy(28)}{dt} = y(28)(-d_7 - k_7) + a_7(y(22)y(27)), \tag{O.28}$$

$$\frac{dy(29)}{dt} = y(29)(-d_7 - k_7) + a_7(y(26)y(27)), \tag{O.29}$$

$$\begin{aligned}\frac{dy(30)}{dt} = {} & y(11)y(30)(-a_8) + d_8y(31) + y(19)y(30)(-a_8) + d_8y(32) \\ & + y(21)y(30)(-a_8) + d_8y(33) + y(23)y(30)(-a_8) + d_8y(34) \\ & + y(24)y(30)(-a_8) + d_8y(35) + y(25)y(30)(-a_8) + d_8y(36),\end{aligned} \tag{O.30}$$

$$\begin{aligned}\frac{dy(31)}{dt} = {} & y(31)(-d_8) + a_8(y(11)y(30)) + y(31)y(2)(-a_5) + d_5y(32) \\ & + y(31)y(20)(-a_6) + d_6y(33) + y(31)y(22)(-a_6) + d_6y(34),\end{aligned} \tag{O.31}$$

$$\begin{aligned}\frac{dy(32)}{dt} = {} & y(32)(-d_8) + a_8(y(19)y(30)) + y(32)(-d_5) + a_5(y(31)y(2)) \\ & + y(32)y(20)(-a_6) + d_6y(35) + y(32)y(22)(-a_6) + d_6y(36),\end{aligned} \tag{O.32}$$

$$\frac{dy(33)}{dt} = y(33)(-d_8) + a_8(y(21)y(30)) + y(33)(-d_6) + a_6(y(31)y(20)) + y(33)y(2)(-a_5) + d_5y(35), \quad (O.33)$$

$$\frac{dy(34)}{dt} = y(34)(-d_8) + a_8(y(23)y(30)) + y(34)(-d_6) + a_6(y(31)y(22)) + y(34)y(2)(-a_5) + d_5y(36), \quad (O.34)$$

$$\frac{dy(35)}{dt} = y(35)(-d_8) + a_8(y(24)y(30)) + y(35)(-d_5) + a_5(y(33)y(2)) + y(35)(-d_6) + a_6(y(32)y(20)), \quad (O.35)$$

$$\frac{dy(36)}{dt} = y(36)(-d_8) + a_8(y(25)y(30)) + y(36)(-d_5) + a_5(y(34)y(2)) + y(36)(-d_6) + a_6(y(32)y(22)), \quad (O.36)$$

$$\frac{dy(37)}{dt} = y(37)(-d_9) + a_9(y(26)y(2)) + y(37)y(38)(-a_{10}) + d_{10}y(40), \quad (O.37)$$

$$\frac{dy(38)}{dt} = y(26)y(38)(-a_{10}) + d_{10}y(39) + y(37)y(38)(-a_{10}) + d_{10}y(40) + y(43)k_{11} + y(45)y(38)(-a_{10}) + d_{10}y(46), \quad (O.38)$$

$$\frac{dy(39)}{dt} = y(39)(-d_{10}) + a_{10}(y(26)y(38)) + y(39)y(2)(-a_9) + d_9y(40) + y(39)y(44)(-a_{12}) + d_{12}y(46), \quad (O.39)$$

$$\frac{dy(40)}{dt} = y(40)(-d_{10} - k_{910}) + a_{10}(y(37)y(38)) + y(40)(-d_9 - k_{910}) + a_9(y(39)y(2)), \quad (O.40)$$

$$\frac{dy(41)}{dt} = y(40)k_{910} + y(40)k_{910} + y(41)y(42)(-a_{11}) + d_{11}y(43), \quad (O.41)$$

$$\frac{dy(42)}{dt} = y(41)y(42)(-a_{11}) + d_{11}y(43) + y(43)k_{11}, \quad (O.42)$$

$$\frac{dy(43)}{dt} = y(43)(-d_{11} - k_{11}) + a_{11}(y(41)y(42)), \quad (O.43)$$

$$\frac{dy(44)}{dt} = y(26)y(44)(-a_{12}) + d_{12}y(45) + y(39)y(44)(-a_{12}) + d_{12}y(46), \quad (O.44)$$

$$\frac{dy(45)}{dt} = y(45)(-d_{12}) + a_{12}(y(26)y(44)) + y(45)y(38)(-a_{10}) + d_{10}y(46), \quad (O.45)$$

$$\frac{dy(46)}{dt} = y(46)(-d_{12}) + a_{12}(y(39)y(44)) + y(46)(-d_{10}) + a_{10}(y(45)y(38)). \quad (O.46)$$

#### 1.3 The conservation laws (C.1-C.11)

The system of ODEs (O.1-O.46) obeys a set of conservation laws (C.1-C.11) which are,

$$0 = y(1) + y(3) + y(5) + y(7) + y(8) + y(10) + y(16) + y(17) + y(18) - BRAFV600E_{tot}, \quad (C.1)$$

$$0 = y(4) + y(5) + y(6) + y(7) + y(8) + y(10) + y(11) + y(13) + y(14) + y(17) + y(18) + y(19) + y(21) + y(23) + y(24) + y(25) + y(31) + y(32) + y(33) + y(34) + y(35) + y(36) - MEK_{tot}, \quad (C.2)$$

$$0 = y(20) + y(21) + y(22) + y(23) + y(24) + y(25) + y(26) + y(28) + y(29) + y(33) + y(34) + y(35) + y(36) + y(37) + y(39) + y(40) + y(45) + y(46) - ERK_{tot}, \quad (C.3)$$

$$0 = y(38) + y(39) + y(40) + y(41) + y(43) + y(46) - S_{tot}, \quad (C.4)$$

$$0 = y(2) + y(3) + y(8) + y(10) + y(19) + y(24) + y(25) + y(32) + y(35) + y(36) + y(37) + y(40) - ATP_{tot}, \quad (C.5)$$

$$0 = y(15) + y(16) + y(17) + y(18) - DBF_{tot}, \quad (C.6)$$

$$0 = y(30) + y(31) + y(32) + y(33) + y(34) + y(35) + y(36) - TMT_{tot}, \quad (C.7)$$
$$0 = y(44) + y(45) + y(46) - SCH_{tot}, \quad (C.8)$$
$$0 = y(12) + y(13) + y(14) - phosph_{1,tot}, \quad (C.9)$$
$$0 = y(27) + y(28) + y(29) - phosph_{2,tot}, \quad (C.10)$$
$$0 = y(42) + y(43) - phosph_{3,tot}. \quad (C.11)$$

Note that we assume an ADP-to-ATP recycling to continually occur in the system, and thus the $[ADP]$ variable is not included in the $[ATP]$ conservation law. This ATP-to-ADP recycling is not explicitly included in the model. In C.1-C.11, the total concentrations, which are denoted by a $tot$-subscript, are equal to the initial conditions used in the simulations so that,

$$\begin{aligned}
BRAFV600E_{tot} &= \text{[BRAFV600E](0)}, \\
MEK_{tot} &= \text{[MEK](0)}, \\
ERK_{tot} &= \text{[ERK](0)}, \\
S_{tot} &= \text{[S](0)}, \\
phosph_{1,tot} &= \text{[phosph_1](0)}, \\
phosph_{2,tot} &= \text{[phosph_2](0)}, \\
phosph_{3,tot} &= \text{[phosph_3](0)}, \\
DBF_{tot} &= \text{[DBF](0)}, \\
TMT_{tot} &= \text{[TMT](0)}, \\
SCH_{tot} &= \text{[SCH](0)}.
\end{aligned}$$

### 2 Supplementary Note 2: Code access and instructions

The computational MATLAB framework that was used to turn the system of reactions (R.1-R.44) into a system of differential-algebraic equations (O.1-O.46, C.1-C.11), and numerically solve these equations to simulate pathway dynamics, was presented in one of our previous articles.[1] Detailed instructions on how to use and manipulate the code files are available in the supplementary material of our previous article (Hamis et al., Br J Cancer, 2021).

The MATLAB code used in the current article is available on GitHub, in the `extended_code/three_drugs` folder in the repository `https://github.com/SJHamis/MAPKcascades`. To tailor the code files used in Hamis et al.[1] for our current system of interest (Fig.1 in the main article), we followed the step-by-step instructions below:

1. Access the original MATLAB codes on `https://github.com/SJHamis/MAPKcascades`.
2. Open the file `extended_code/three_drugs/model_details/set_system_of_reactions.m` and modify the system of reactions to describe the model of interest.
3. Open the file `extended_code/three_drugs/model_details/set_reaction_constants.m` and modify the rate constants to describe the rate constants of interest.
4. Run the file `extended_code/three_drugs/auxiliary_files_model_setup/main_setup_convert_SoRs_to_SoODEs.m`. This will overwrite the file `extended_code/three_drugs/auxiliary_files_model_setup/mapk_cascade_DAE.m` to describe the newly implemented model. The auxiliary file `extended_code/three_drugs/auxiliary_files_model_setup/get_conslaw_position.m`, which is used to include conservation laws in the DAE, will also be updated.
5. Use the MATLAB files that start with the keyword `main` to generate numerical simulations and result-figures. Edit the main files to plot the output(s) of interest.

### 3 Supplementary Note 3: Data and estimation of model parameters

#### 3.1 Rate constants

The rate constants in the system of reactions (R.1-R.44), and the system of ODEs (O.1-O.46) are listed in Supplementary Table 1. A discussion on how the constants pertaining to reactions R.1-R.36 are estimated from in vitro data is available in the Supplementary Material of Hamis et al..[1] A discussion on how the constants pertaining to reactions R.37-R.44 are estimated from in vitro data is available in Section 3.3 of this document.

Supplementary Table 1: **Rate constants in the system of reactions (R.1-R.44).** The system of reactions includes 12 forward rate constants $a_1, a_2, .., a_{12}$, 12 reverse rate constants $d_1, d_2, .., d_12$, and 6 catalytic rate constants $k_{1,2}, k_3, k_{5,6}, k_7, k_{9,10}, k_{11}$. In the description column, stroke symbols (/) denote 'or', and ampersand symbols (&) denote 'and'. The rightmost column lists the chemical reactions (SM1.1) in which the constants appear. † All forward rate constants have the same value.

| Rate constants in the system of reactions (R.1-R.44) | | | | |
|---|---|---|---|---|
| constant | value | reference | description | appearance |
| $a_1$ | 0.106$\mu$M$^{-1}$s$^{-1}$ | [2, 3] | Forward rate constant for ATP binding to BRAFV600E (or a BRAFV600E complex with MEK/pMEK). | R.1,R.6,R.7 |
| $d_1$ | 6.23 s$^{-1}$ | [2, 3, 4] | Reverse rate constant for ATP binding to BRAFV600E (or a BRAFV600E complex with MEK/pMEK). | R.1,R.6,R.7 |
| $a_2$ | 0.106$\mu$M$^{-1}$s$^{-1}$ | † | Forward rate constant for MEK/pMEK binding to BRAFV600E (or a BRAFV600E complex with ATP/DBF). | R.2,R.3,R.4, R.5,R.13,R.14 |
| $d_2$ | 0.02385 s$^{-1}$ | [2, 3] | Reverse rate constant for MEK/pMEK binding to BRAFV600E (or a BRAFV600E complex with ATP/DBF). | R.2,R.3,R.4, R.5,R.13,R.14 |
| $k_{1,2}$ | 0.66 s$^{-1}$ | [3] | Catalytic rate constant for production of pMEK/ppMEK from BRAFV600E·ATP complex with MEK/pMEK. | R.4,R.5,R.6, R.7 |
| $a_3$ | 0.106$\mu$M$^{-1}$s$^{-1}$ | † | Forward rate constant for phosph$_1$ binding to pMEK/ppMEK. | R.8,R.9 |
| $d_3$ | 0.0159 s$^{-1}$ | [2, 3] | Reverse rate constant for phosph$_1$ binding to pMEK/ppMEK. | R.8,R.9 |
| $k_3$ | 0.0159 s$^{-1}$ | [2, 3] | Catalytic rate constant for production of MEK/pMEK from phosph$_1$ complex with pMEK/ppMEK. | R.8,R.9 |
| $a_4$ | 0.106$\mu$M$^{-1}$s$^{-1}$ | † | Forward rate constant for the drug DBF binding to BRAFV600E (or a BRAFV600E complex with MEK/pMEK). | R.10,R.11,R.12 |
| $d_4$ | 0.0000593 s$^{-1}$ | [2, 3, 5, 6, 7, 8] | Reverse rate constant for the drug DBF binding to BRAFV600E (or a BRAFV600E complex with MEK/pMEK). | R.10,R.11,R.12 |
| $a_5$ | 0.106$\mu$M$^{-1}$s$^{-1}$ | † | Forward rate constant for ATP binding to ppMEK (or a ppMEK complex with ERK/pERK/TMT/TMT&ERK/TMT&pERK). | R.15,R.20,R.21, R.30,R.33,R.34 |
| $d_5$ | 0.3468 s$^{-1}$ | [2, 3, 9] | Reverse rate constant for ATP binding to ppMEK (or a ppMEK complex with ERK/pERK/TMT/TMT&ERK/TMT&pERK). | R.15,R.20,R.21, R.30,R.33,R.34 |
| $a_6$ | 0.106$\mu$M$^{-1}$s$^{-1}$ | † | Forward rate constant for ERK/pERK binding to ppMEK (or a ppMEK complex with ATP/TMT/TMT&ATP). | R.16,R.17,R.18, R.19,R.31,R.32, R.35,R.36 |

Supplementary Table 1: **Rate constants in the system of reactions (R.1-R.44).** The system of reactions includes 12 forward rate constants $a_1, a_2, .., a_{12}$, 12 reverse rate constants $d_1, d_2, .., d_12$, and 6 catalytic rate constants $k_{1,2}, k_3, k_{5,6}, k_7, k_{9,10}, k_{11}$. In the description column, stroke symbols (/) denote 'or', and ampersand symbols (&) denote 'and'. The rightmost column lists the chemical reactions (SM1.1) in which the constants appear. † All forward rate constants have the same value.

| Rate constants in the system of reactions (R.1-R.44) | | | | |
|---|---|---|---|---|
| constant | value | reference | description | appearance |
| $d_6$ | 0.01184 $s^{-1}$ | [2, 3, 9] | Reverse rate constant for ERK/pERK binding to ppMEK (or a ppMEK complex with ATP/TMT/TMT&ATP). | R.16,R.17,R.18, R.19,R.31,R.32, R.35,R.36 |
| $k_{5,6}$ | 0.0242 $s^{-1}$ | [9] | Catalytic rate constant for production of pERK/ppERK from ppMEK·ATP complex with ERK/pERK. | R.18,R.19,R.20, R.21 |
| $a_7$ | 0.106$\mu M^{-1}s^{-1}$ | † | Forward rate constant for $phosph_2$ binding to pERK/ppERK. | R.22,R.23 |
| $d_7$ | 0.0159 $s^{-1}$ | [2, 3] | Reverse rate constant for $phosph_2$ binding to pERK/ppERK. | R.22,R.23 |
| $k_7$ | 0.0159 $s^{-1}$ | [2, 3] | Catalytic rate constant for production of ERK/pERK from $phosph_2$ complex with pERK/ppERK. | R.22,R.23 |
| $a_8$ | 0.106$\mu M^{-1}s^{-1}$ | † | Forward rate constant for the drug TMT binding to ppMEK (or a ppMEK complex with ATP/ERK/pERK/ERK&ATP/pERK&ATP). | R.24,R.25,R.26, R.27,R.28,R.29 |
| $d_8$ | 0.0012296 $s^{-1}$ | [2, 3, 10] | Reverse rate constant for the drug TMT binding to ppMEK (or a ppMEK complex with ATP/ERK/pERK/ERK&ATP/pERK&ATP). | R.24,R.25,R.26, R.27,R.28,R.29 |
| $a_9$ | 0.106$\mu M^{-1}s^{-1}$ | [2, 3] | Forward rate constant for ATP binding to ppERK (or a ppERK complex with S). | R.37,R.40 |
| $d_9$ | 14.84 $s^{-1}$ | [11, 1, 2] | Reverse rate constant for ATP binding to ppERK (or a ppERK complex with S). | R.37,R.40 |
| $a_{10}$ | 0.106$\mu M^{-1}s^{-1}$ | † | Forward rate constant for S binding to ppERK (or a ppERK complex with ATP/SCH). | R.38,R.39,R.44 |
| $d_{10}$ | 0.000421 $s^{-1}$ | [11, 2, 12] | Reverse rate constant for S binding to ppERK (or a ppERK complex with ATP/SCH). | R.38,R.39,R.44 |
| $k_{9,10}$ | 0.000639 $s^{-1}$ | [11] | Catalytic rate constant for production of pS from ppERK·ATP complex with S/pS. | R.39,R.40 |
| $a_{11}$ | 0.106$\mu M^{-1}s^{-1}$ | † | Forward rate constant for $phosph_3$ binding to pS. | R.41 |
| $d_{11}$ | 0.0159 $s^{-1}$ | [1, 2] | Reverse rate constant for $phosph_3$ binding to pS. | R.41 |
| $k_{11}$ | 0.0159 $s^{-1}$ | [1, 2] | Catalytic rate constant for production of S from $phosph_3$ complex with pS/ppS. | R.41 |
| $a_{12}$ | 0.106$\mu M^{-1}s^{-1}$ | † | Forward rate constant for the drug SCH binding to ppERK (or a ppERK complex with S). | R.42,R.43 |
| $d_{12}$ | 0.00022967 $s^{-1}$ | [1, 2, 13] | Reverse rate constant for the drug SCH binding to ppERK (or a ppERK complex with S). | R.42,R.43 |

#### 3.2 Total and initial concentrations of pathway components

The initial concentrations of signalling molecules that were used in all simulations (Figs. 2-4 in the main manuscript) are listed in Supplementary Table 2.

| Initial concentrations | | |
|---|---|---|
| initial condition | value | reference |
| [BRAFV600E](0) | 3-10 nM* | [2] |
| [MEK](0) | 1.2 $\mu$M | [2] |
| [ERK](0) | 1.2 $\mu$M | [2] |
| [S](0) | 1.2 $\mu$M | [2] |
| [phosph1](0) | 0.3 nM | [2] |
| [phosph2](0) | 0.12 $\mu$M | [2] |
| [phosph3](0) | 0.12 $\mu$M | [2] |
| [ATP](0) | 1-5 mM** | [9] |
| [DBF](0) | *applied DBF dose* (here 0-10 $\mu$M) | |
| [TMT](0) | *applied TMT dose* (here 0-10 $\mu$M) | |
| [SCH](0) | *applied SCH dose* (here 0-10 $\mu$M) | |

Supplementary Table 2: Model initial conditions, where the (0) notation denotes 'at time zero'. For molecules not listed in this table, the initial conditions are zero. *Huang & Ferrell[2] listed MAPKKK = 3 nM, and we use this as our baseline value. However, in order to study amplified BRAFV600E levels as a mode of drug resistance, we explore BRAFV600E initial condition values within the range of 3-10 nM. Similarly, ATP concentrations are explored within the range of 1-5 mM, but the baseline calibrated value is 1 mM.

#### 3.3 Estimating rate constants for reactions downstream of activated ERK (ppERK)

##### 3.3.1 Estimating rate constants for ERK-ATP reactions

In the system of reactions (R.1-R.44), $a_9$, $d_9$ and $k_{9,10}$ respectively denote the association (forward), dissociation (reverse), and catalytic rate constants for ERK-ATP reactions (including pERK-ATP and ppERK-ATP reactions). Lee et al.[11] reported that the catalytic rate constant for a substrate's phosphorylation (the transcription factor Myc) is $k_{cat} = 2.3hr^{-1} \approx 6.39 \times 10^{-4}\ s^{-1}$. Moreover, Callaway et al.[14] reported that the Michaelis constant for ERK-ATP reactions is $K_m^{ATP} = (140.0 \pm 5.0)\ \mu M \approx 140.0\ \mu M$. We first use these data to directly estimate $k_{9,10} = k_{cat} = 6.39 \times 10^{-4}\ s^{-1}$. To next derive values for $a_9$ and $d_9$, we note that Huang and Ferrell[2] argued that it is the Michaelis constants, $K_{mi} = (d_i + k_i)/a_i$, that are of importance when computing steady-state enzyme levels, rather than the absolute values of $d_i, a_i, k_i$. Following this argument, we set $a_9 = a_1 =$0.106 $s^{-1}\,(\mu M)^{-1}$ and calculate $d_9$ from Lee et al.[11] and Callaway et al.'s[14] data. By doing this, we obtain

$$
\begin{aligned}
& K_m^{ATP} = \frac{d_9 + k_{9,10}}{a_9} \\
\rightarrow\ & a_9 \cdot K_m^{ATP} = d_9 + k_{9,10} \\
\rightarrow\ & d_9 = a_9 \cdot K_m^{ATP} -\ k_{9,10} \\
\rightarrow\ & d_9 \approx 0.106\ s^{-1}\,(\mu M)^{-1} \cdot 140.0\ \mu M - 6.39 \times 10^{-4}\ s^{-1} \approx 14.84\ s^{-1}\ .
\end{aligned}
\tag{P.1}
$$

Note that all forward rate constants in the system are the same (Supplementary Table 1), following Huang and Ferrell.[2]

##### 3.3.2 Estimating rate constants for ERK-substrate interactions

In the system of reactions (R.1-R.44), $a_{10}$, and $d_{10}$ respectively denote the association (forward) and dissociation (reverse) rate constants for ppERK-S reactions (including ppERK-pS reactions).

Lee et al.[11], reported that the Michaelis-Menten constant for a substrate's phosphorylation (the transcription factor Myc) is $K_m^{MYC} = 0.01\mu M$. Thus, we can obtain $d_{10}$ from the $K_m^{MYC}$ data using

$$\begin{aligned} K_m^{MYC} &= \frac{d_{10} + k_{9,10}}{a_{10}} \\ \rightarrow\ d_{10} &= a_{10} \cdot K_m^{MYC} -\ k_{9,10} \\ \rightarrow\ d_{10} &\approx 0.106\ s^{-1}\,(\mu M)^{-1} \cdot 0.01\mu M - 6.39 \times 10^{-4} s^{-1} \approx 4.21 \times 10^{-4} s^{-1}, \end{aligned} \tag{P.2}$$

following a relationship between binding constants explained in Motulsky and Neubig.[12]

##### 3.3.3 Estimating rate constants for substrate-phosphatase interactions

In the system of reactions (R.1-R.44), $a_{11}$, $d_{11}$ and $k_{11}$ respectively denote the association (forward), dissociation (reverse) and catalytic rate constants for pS-phosph$_3$ reactions. We first set $a_{11}$ = $a_1$ =0.106 $s^{-1}\,(\mu M)^{-1}$, following the discussion in Section 3.3.1. We next note that Huang and Ferrell[2] estimated the Michaelis constant $K_m$ to be $0.3\mu M$ for chemical reactions in their regarded MAPKKK-MAPKK-MAPK system. To enable an estimation of $d_{11}$, we assume that this assumption holds for phosph$_3$ reactions in our BRAFV600E-MEK-ERK system (which is based on Huang and Ferrell's[2] model). For simplicity, we also assume that $d_{11} = k_{11}$,[1] and obtain

$$\begin{aligned} K_m &= \frac{d_{11} + k_{11}}{a_{11}} \\ \rightarrow\ d_{11} + k_{11} &= K_m \cdot a_{11} \\ \rightarrow\ \text{(with } d_{11} = k_{11}\text{)}\ \rightarrow d_{11} &= k_{11} = \frac{K_m \cdot a_{11}}{2} \\ \rightarrow\ d_{11} = k_{11} &\approx \frac{0.3\ \mu M \cdot 0.106\ s^{-1}\,(\mu M)^{-1}}{2} \approx 0.0159\ s^{-1}\ . \end{aligned} \tag{P.3}$$

##### 3.3.4 Estimating rate constants for ppERK-SCH interactions

In the system of reactions (R.1-R.44), $a_{12}$ and $d_{12}$ respectively denote the association (forward) and dissociation (reverse) rate constants for ppERK-SCH interactions. We first set $a_{12} = a_1$ =0.106 $s^{-1}\,(\mu M)^{-1}$, following the discussion in Section 3.3.1. We next use data pertaining to ppERK-SCH interactions reported by Morris et al.[13] to estimate $d_{12}$.

Morris et al.[13] reported that the IC$_{50}$ values, respectively, are 4 nM/L and 1 nM/L for SCH772984 inhibition of ERK1 and ERK2 activity *in vitro*. From these data, we use the Cheng-Prusoff[15,16] approach to estimate the inhibitory constant $K_i$ as

$$K_i = \frac{IC_{50}}{1 + [L]\ /\ K_d} = \frac{(IC_{50,ERK1} + IC_{50,ERK2})\,/2}{1 + [L]\ /\ K_d}, \tag{P.4}$$

where $[L]$ denotes the concentration of the experimental ligand (ATP), and $K_d$ is the equilibrium dissociation constant between ATP and ppERK. Approximang $[L] \approx 10\mu M$,[1] and $K_d \approx K_m = 65\mu M$,[1] we obtain

$$K_i \approx \frac{(4+1)/2\ nM}{1 + 10\ \mu M\ /\ 65\ \mu M} \approx 2.17\ nM. \tag{P.5}$$

We can now estimate $d_{12}$ from $K_i$,[12]

$$K_{i,12} = \frac{d_{12}}{a_{12}} \rightarrow d_{12} \approx 2.17\ nM \cdot 0.106\ s^{-1}\,(\mu M)^{-1} \approx 2.2967 \times 10^{-4}\ s^{-1}\,. \qquad \text{(P.6)}$$

### 4 Supplementary Note 4: Demonstrating the transient behaviour of system dynamics in response to SCH monotherapies

Supplementary Fig.1 demonstrates the transient behaviour of system dynamics in response to SCH monotherapies for baseline and elevated BRAFV600E concentrations.

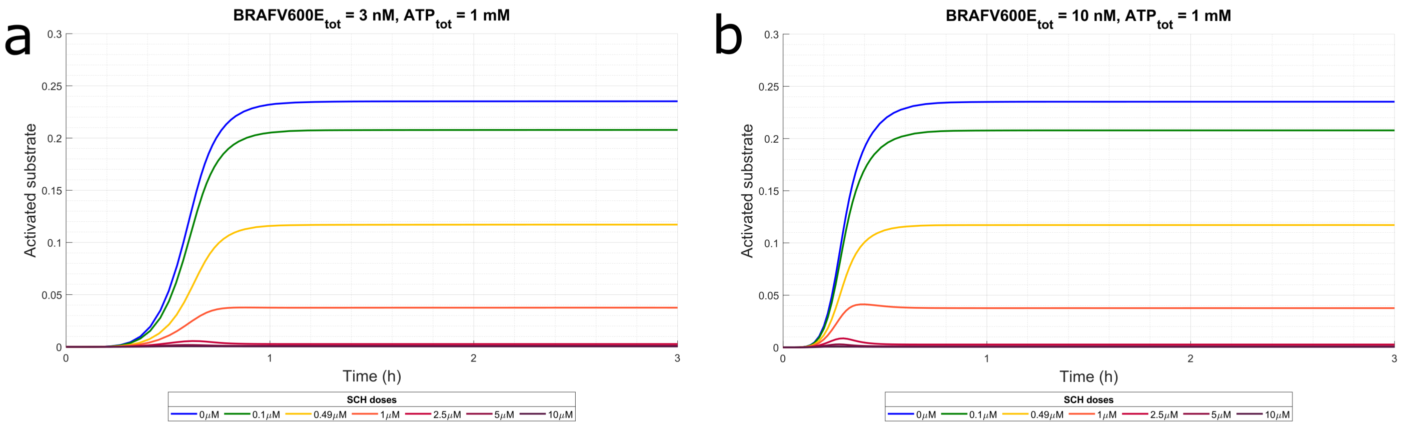

Supplementary Fig. 1: **The system displays transient behaviour in response to SCH monotherapies**. Substrate activity levels ($S_{\text{act}}(t)$) are plotted over time for various SCH monotherapies in the range 0 $\mu$M to 10 $\mu$M. The baseline value for total ATP (1 mM) concentration is used. (a) The simulations are produced using the estimated model parameters listed in Supplementary Table 1, for BRAFV600E = 3 nM concentration. The SCH dose $0.49\mu$M inhibits 50% of the substrate activity at the steady state. (b) The BRAFV600E concentration has been raised to 10 nM.

### 5 Supplementary Note 5: Identifying two-drug combinations that yield synergistic responses

Supplementary Fig.2 is a binary reconstruction of Fig.3 in the main manuscript. Isoboles that connect the monotherapies that yield 90% inhibition are included in the right-most column. Drug combinations under the isoboles that yield 90% substrate inhibition or more are here classified as synergistic.

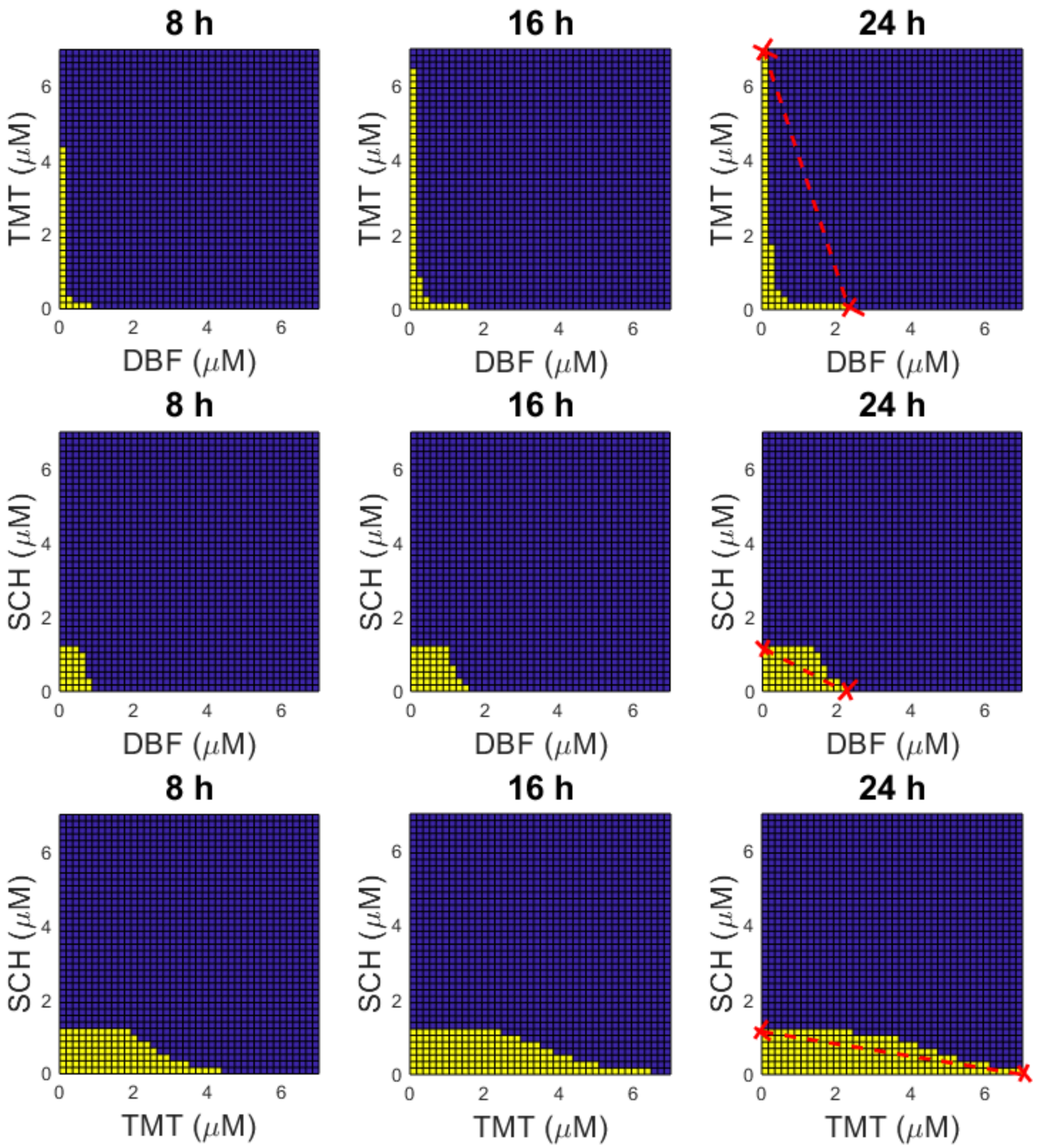

Supplementary Fig. 2: Drug dose combinations that yield a substrate activation of 90% or more are indicated by blue bins in the heatmaps. Other dose combinations are indicated by yellow bins.

### 6 Supplementary Note 6: Identifying three-component vertical inhibition strategies that achieve desired substrate activation levels

In the main manuscript, we used the mathematical model (Fig.1 in the main manuscript) to identify three-component vertical inhibition strategies that achieve 90% inhibition of substrate activation (such that $S^*_{act}(t) = 0.1$) at selected time points (Fig.4 in the main manuscript). To aid in reading this figure, we provide a data matrix from which DBF, TMT, and SCH doses that achieve 90%

inhibition of substrate activation at 24 hours can be read numerically (Supplementary Fig.3). The matrix shows that, for instance, three (DBF,TMT,SCH) doses that achieve $S_{act}(\text{24 hours}) \leq 0.1$ are (0,0,1.2 $\mu$M), (0.3 $\mu$M,0.4 $\mu$M,0.75 $\mu$M), and (0.1 $\mu$M,2 $\mu$M,0.6 $\mu$M).

To further exemplify that the model can be used to find other, user-defined, substrate activity levels, we here use the model to identify strategies that achieve 75% (Supplementary Fig.4), 50% (Supplementary Fig.5), and 25% (Supplementary Fig.6) inhibition of substrate activity at 8, 16, and 24 hours. In the figures, dose combinations that lie under the red synergy planes are identified as synergistic.

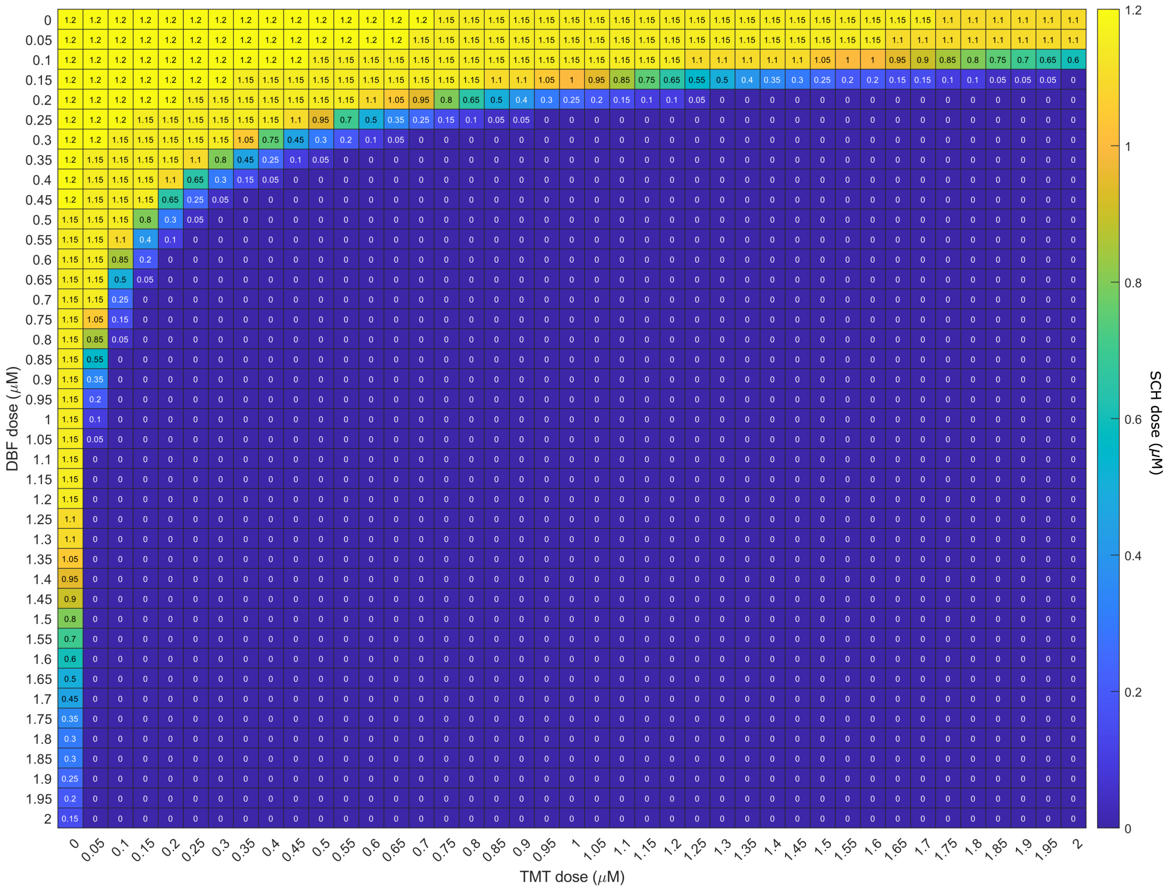

Supplementary Fig. 3: **Drug-dose combinations that achieve 90% inhibition of substrate activation at 24 hours for baseline BRAV600E and ATP concentrations can be read from a data matrix.** The data matrix is used to produce the drug-dose combination surface in the the top-right plot of Fig.4 in the main manuscript. Each row in the matrix corresponds to a DBF dose, and each column in the matrix corresponds to a TMT dose. For each DBF dose x and TMT dose y, the numerical value z in the matrix element corresponds to the SCH dose needed to achieve inhibition of 90% substrate activity, given x and y.

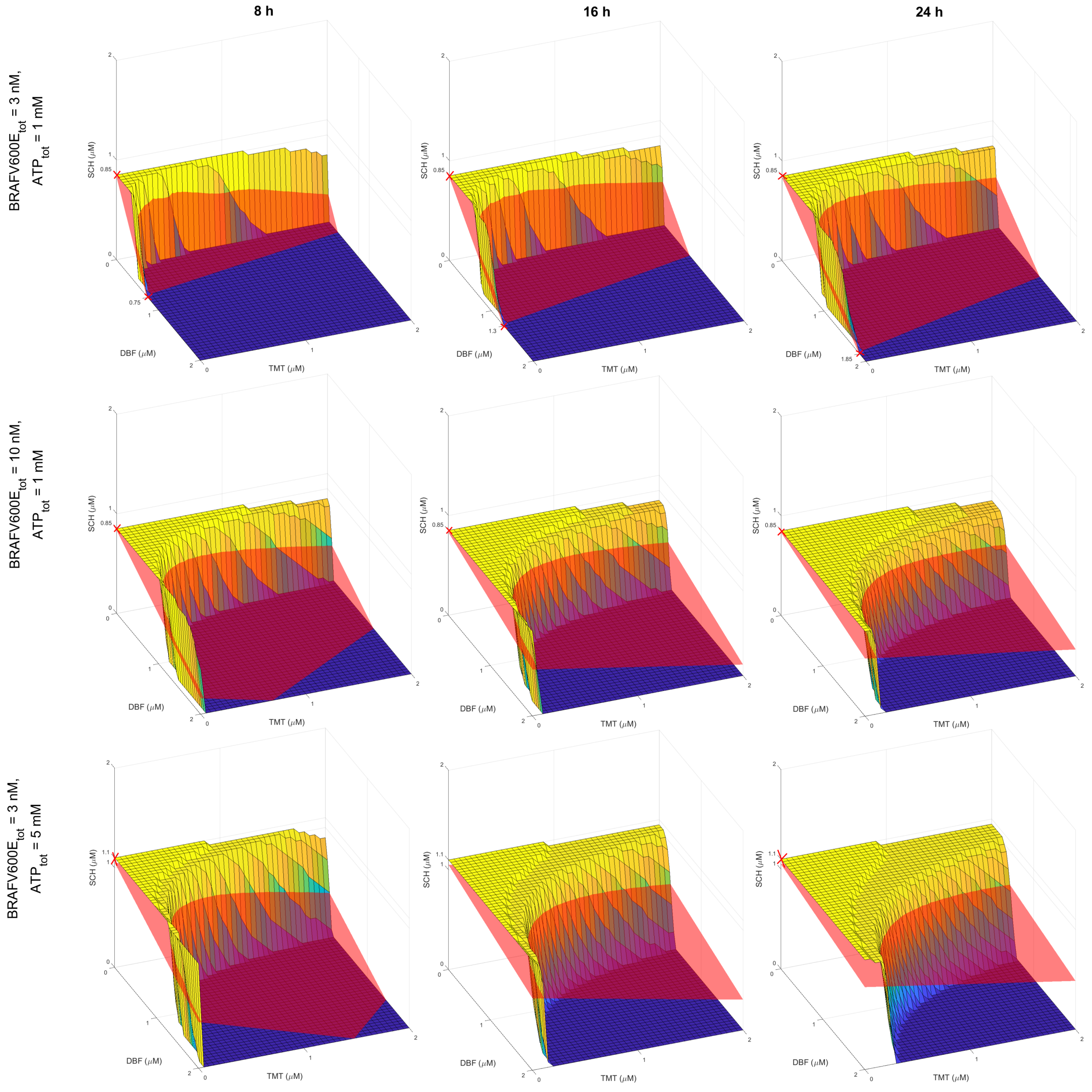

Supplementary Fig. 4: **Mathematical modelling quantifies *in silico* treatment responses to vertical inhibition treatment strategies that target up to three components of the BRAFV600E-MEK-ERK pathway and inhibit 75% substrate activity.** The yellow-to-blue surface plots show DBF-TMT-SCH combination therapies that yield 75% substrate activation (so that $S^*_{act}(t) = 0.25$, Eq.3 in the main article). The red synergy-isoboles (planes) connect the monotherapy doses that yield a 75% inhibition of substrate activity. Monotherapy doses that yield 75% substrate activity are marked with crosses. Treatment combinations, *i.e.,* parts of the $S^*_{act}(t) = 0.25$ surfaces, that lie under the synergy-isoboles are here categorised as synergistic. Results are plotted at 8 (left column), 16 (middle column), and 24 (right column) hours for baseline total ATP and BRAFV600E concentrations (top row), and for elevated BRAFV600E concentrations (middle row) and ATP concentrations (bottom row).

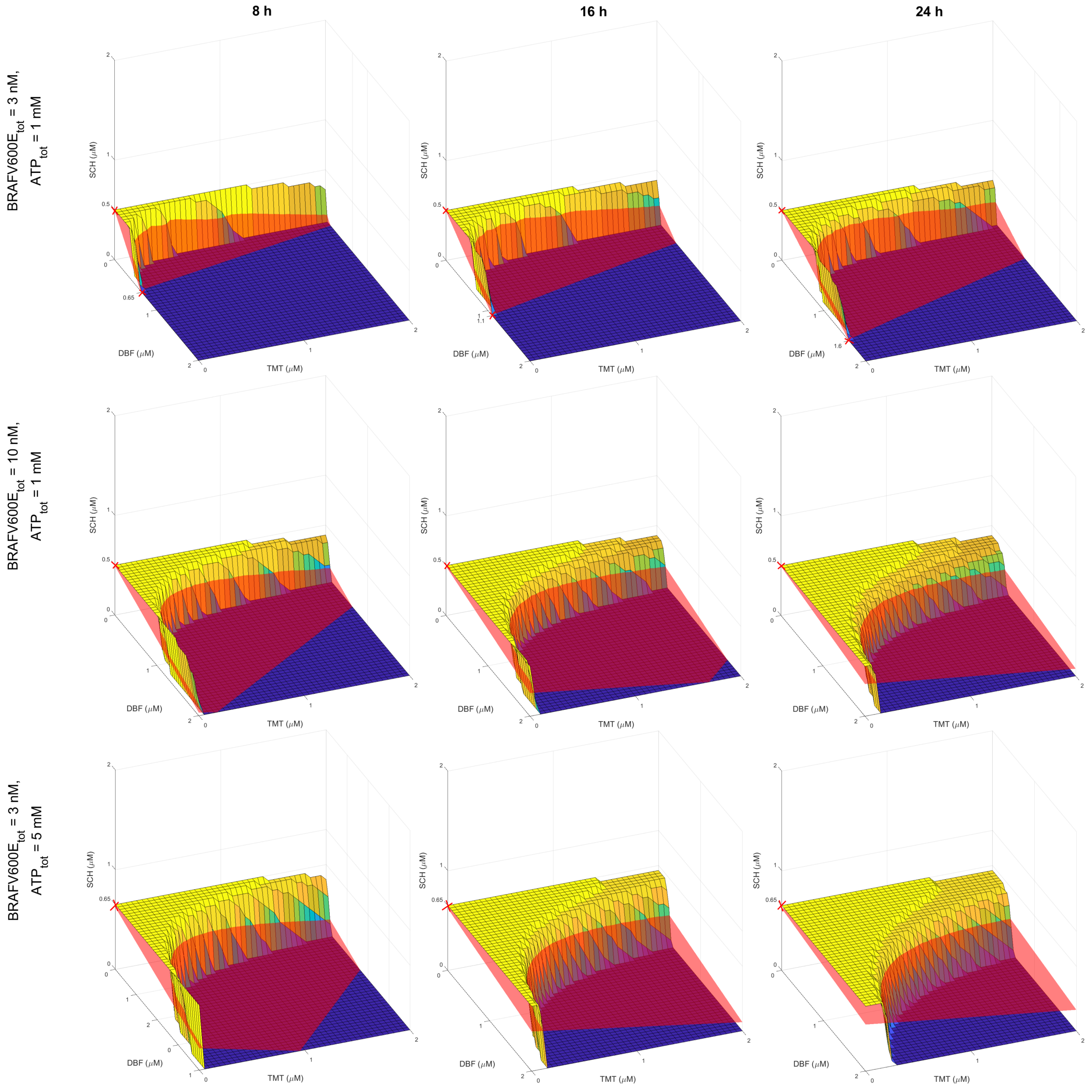

Supplementary Fig. 5: **Mathematical modelling quantifies *in silico* treatment responses to vertical inhibition treatment strategies that target up to three components of the BRAFV600E-MEK-ERK pathway and inhibit 50% substrate activity.** The yellow-to-blue surface plots show DBF-TMT-SCH combination therapies that yield 50% substrate activation (so that $S^*_{act}(t) = 0.5$, Eq.3 in the main article). The red synergy-isoboles (planes) connect the monotherapy doses that yield a 50% inhibition of substrate activity. Monotherapy doses that yield 50% substrate activity are marked with crosses. Treatment combinations, *i.e.,* parts of the $S^*_{act}(t) = 0.5$ surfaces, that lie under the synergy-isoboles are here categorised as synergistic. Results are plotted at 8 (left column), 16 (middle column), and 24 (right column) hours for baseline total ATP and BRAFV600E concentrations (top row), and for elevated BRAFV600E concentrations (middle row) and ATP concentrations (bottom row).

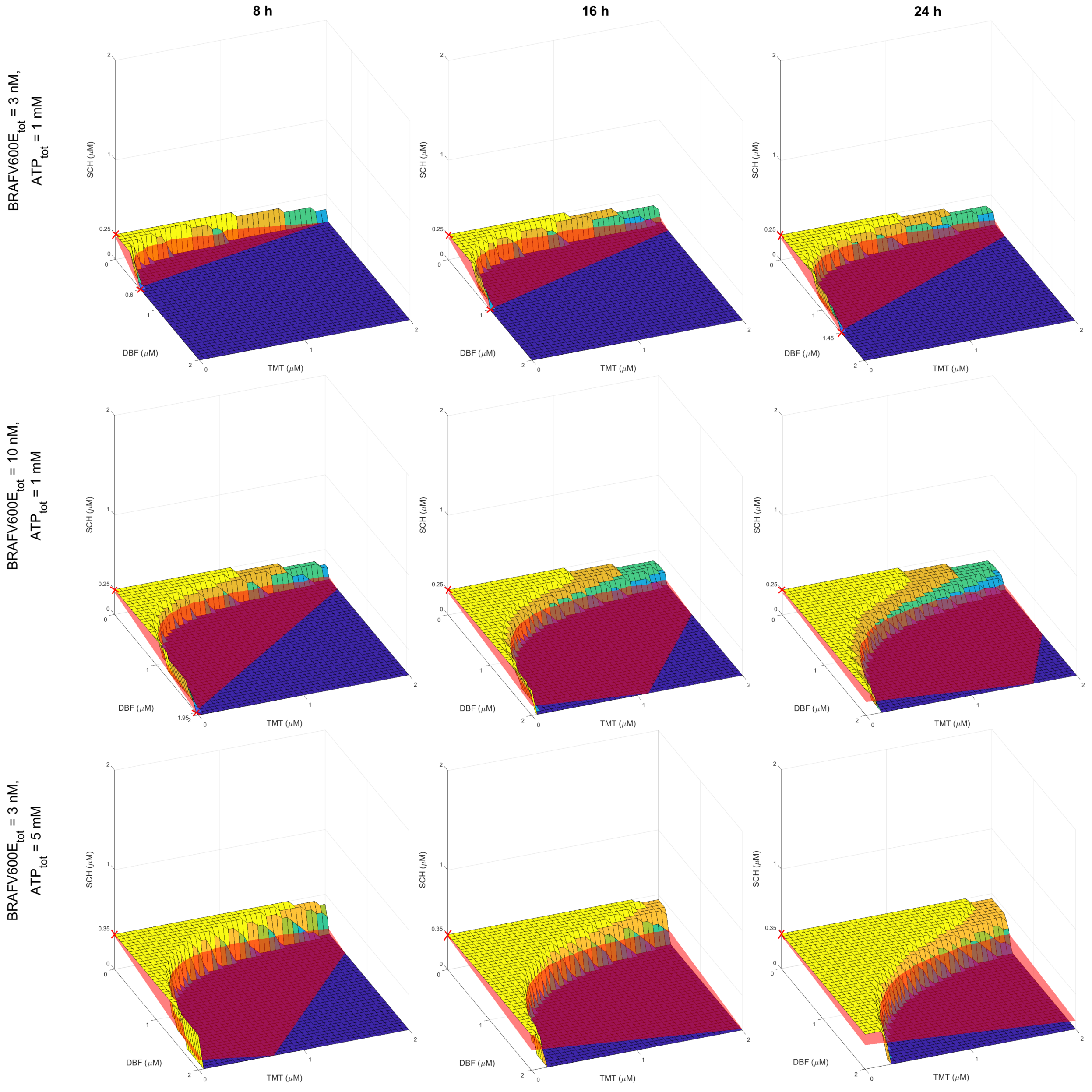

Supplementary Fig. 6: **Mathematical modelling quantifies *in silico* treatment responses to vertical inhibition treatment strategies that target up to three components of the BRAFV600E-MEK-ERK pathway and inhibit 25% substrate activity.** The yellow-to-blue surface plots show DBF-TMT-SCH combination therapies that yield 25% substrate activation (so that $S^*_{act}(t) = 0.75$, Eq.3 in the main article). The red synergy-isoboles (planes) connect the monotherapy doses that yield a 25% inhibition of substrate activity. Monotherapy doses that yield 25% substrate activity are marked with crosses. Treatment combinations, *i.e.,* parts of the $S^*_{act}(t)$ =0.75 surfaces, that lie under the synergy-isoboles are here categorised as synergistic. Results are plotted at 8 (left column), 16 (middle column), and 24 (right column) hours for baseline total ATP and BRAFV600E concentrations (top row), and for elevated BRAFV600E concentrations (middle row) and ATP concentrations (bottom row).
